# Co-grafting strategies uncover cell type–dependent regulation of dopamine neuron specification and functional maturation in a pre-clinical model of Parkinson’s Disease

**DOI:** 10.64898/2025.12.19.695615

**Authors:** Edoardo Sozzi, María García Garrote, Janitha Mudannayake, German Ramos Passarello, Andreas Bruzelius, Sara Corsi, Greta Galeotti, Mette Habekost, Linda Scaramuzza, Janko Kajtez, Dario Besusso, Anders Björklund, Elena Cattaneo, Alessandro Fiorenzano, Petter Storm, Malin Parmar

## Abstract

Parkinson’s disease (PD), the second most common neurodegenerative disorder, is characterized by the progressive loss of A9 dopaminergic neurons in the *substantia nigra*, leading to dopamine (DA) depletion in the striatum and subsequent motor symptoms. Transplantation of ventral midbrain-patterned DA (vmDA) progenitors derived from human pluripotent stem cells, aimed at restoring DA neurotransmission in the striatum, is being developed and currently explored in ongoing clinical trials. One factor that may improve the maturation and fate determination of DA neurons *in vivo* is the intercellular communication within the graft environment, ultimately affecting the therapeutic outcome. In this study, we co-transplanted vmDA progenitors with either glial, ventral forebrain or striatal progenitors into a preclinical xenograft PD model to investigate how these interactions shape the development, maturation, and function of therapeutic DA neurons. Our findings show that co-grafts with ventral forebrain progenitors increase the yield of DA neurons and also promote their functional maturation. Furthermore, we demonstrated that co-grafts with striatal neurons promote functional maturation and the acquisition of DA subtype identity. From these data, we identified *EBF3* and *PBX3* as candidate transcription factors directing DA neuron maturation and subtype specification, and then functionally validated their role in brain organoids. Taken together, our data highlight that the cellular microenvironment, including specific interactions with neighbouring cells, guides *in vivo* DA neuron specification and maturation. These findings provide a foundation for developing more refined and effective cell preparations for replacement therapy in PD, and define a conceptual framework that could inform stem cell–based strategies for other neurodegenerative diseases.

## Introduction

Parkinson’s disease (PD) is a debilitating neurodegenerative disorder characterized by the progressive loss of dopamine (DA) neurons located in the ventral midbrain (VM), leading to DA depletion in the striatum and the emergence of motor symptoms such as bradykinesia, tremor, rigidity, and postural instability^1^. Current treatments provide symptomatic relief but do not address the underlying neuronal loss, prompting the development of better therapies^1,2^. Transplantation of VM-patterned DA progenitors (vmDA) derived from human pluripotent stem cells is a promising regenerative approach for the treatment of PD, aiming to replace the lost DA neurons and restore physiological levels of DA in the striatum^3,4^, and is being evaluated in multiple ongoing clinical trials^5–14^.

Leading up to these clinical trials, the therapeutic potential of transplanted DA neurons has been extensively investigated and characterized in xenograft models using both fetal and stem cell-derived DA progenitors^15–18^. Recent single-cell analyses have uncovered extensive molecular heterogeneity within these grafts, encompassing both neuronal and non-neuronal cell types^19–23^. We recently showed that the graft environment is likely to play a role in guiding the fate commitment and terminal differentiation of mature cell types within the graft^20^. However, although grafts are known to be heterogeneous in cellular composition, the role of non-DA cells within the graft environment in shaping DA neuron development and function remains largely unexplored.

Another prompt to address this question emerged while performing the dose-finding studies for the clinical vmDA product STEM-PD^5^. In these studies, we diluted a progressively decreasing number of vmDA progenitors with an increasing number of non-DA carrier cells of ventral forebrain (vFB) identity to retain the same density and volume of cell preparation in all grafts. We found that co-grafts containing lower number of STEM-PD progenitors diluted with vFB carrier cells resulted in a comparable number of DA neurons in the mature grafts 6 months post-transplant compared to transplants with same dose of STEM-PD cells alone. This unexpected finding suggests that non-DA carrier cells may enhance DA neuron survival or maturation, supporting the hypothesis that the cellular environment surrounding transplanted DA neurons impacts their lineage commitment, specification, and function which inspired the design of new studies to investigate how support cells influence DA graft development.

In this study, we therefore systematically explored the impact of defined co-graft environments on DA neuron lineage commitment, specification and function. We assessed the influence of three co-grafted cell types: the same vFB cells as used in the pre-clinical validation of STEM-PD^5^; glial progenitors (GPCs), known to support neuronal survival and differentiation both *in vitro* and *in vivo*^25,26^; and STR progenitors that give rise to striatal medium spiny neurons (MSNs), the primary postsynaptic targets of the A9 subtype of DA neurons. We confirmed earlier findings from the efficacy study^5^ showing that co-grafting vmDA progenitors with vFB progenitors markedly increases the yield of DA neurons after grafting, and here demonstrate that this also leads to faster motor recovery, indicating that a specific graft environment could enhance the efficacy of cell replacement therapy for PD. Furthermore, single-cell transcriptional profiling of co-grafts revealed diverse neuronal subtypes, including vmDA-derived DA neurons with molecular signatures corresponding to mature A9 and A10-like subtype identities in different proportions depending on the cellular context of the graft. Interestingly, while co-grafts containing striatal MSNs did not increase the total number of TH neurons, they selectively increased the A9 component of the graft, which also correlated with faster motor recovery.

Gene regulatory network analysis and *in silico* perturbations using CellOracle^27^ identified *EBF3* and *PBX3* as transcriptional regulators of DA neuron specification and maturation. Functional validation in VM organoids demonstrated that both *EBF3* and *PBX3* promote a transcriptional shift toward a more mature DA neuron state, with PBX3 enhancing spontaneous activity and DA release. These findings support a model in which target- or cell-dependent cues act, at least in part, through transcriptional regulators such as *PBX3* and *EBF3* to drive the maturation and subtype specification of stem cell-derived human DA neurons developing within a transplant environment.

## Results

### Co-grafting vmDA progenitors with vFB progenitors results in a higher yield of DA neurons

To assess the impact of non-DA cell types on DA neuron survival, function and maturation, we designed a series of co-grafting experiments where vmDA progenitors were co-transplanted with different cell types to evaluate their effect on subtype-specific differentiation, maturation and function of the DA neurons. In these experiments, we co-grafted vmDA progenitors with GPCs, vFB and STR as illustrated in Fig. 1A. All grafted cells were generated from human embryonic stem cells (hESCs) following established protocols^28–30^ and correct regional patterning was confirmed by gene and protein expression (Suppl. Fig. 1A-K). To track the progeny of each cell type within the graft, all four progenitor populations were labelled *in□vitro* with distinct, heritable GFP-containing genomic barcode libraries prior to transplantation (see Material and Methods for details).

**Figure 1.**
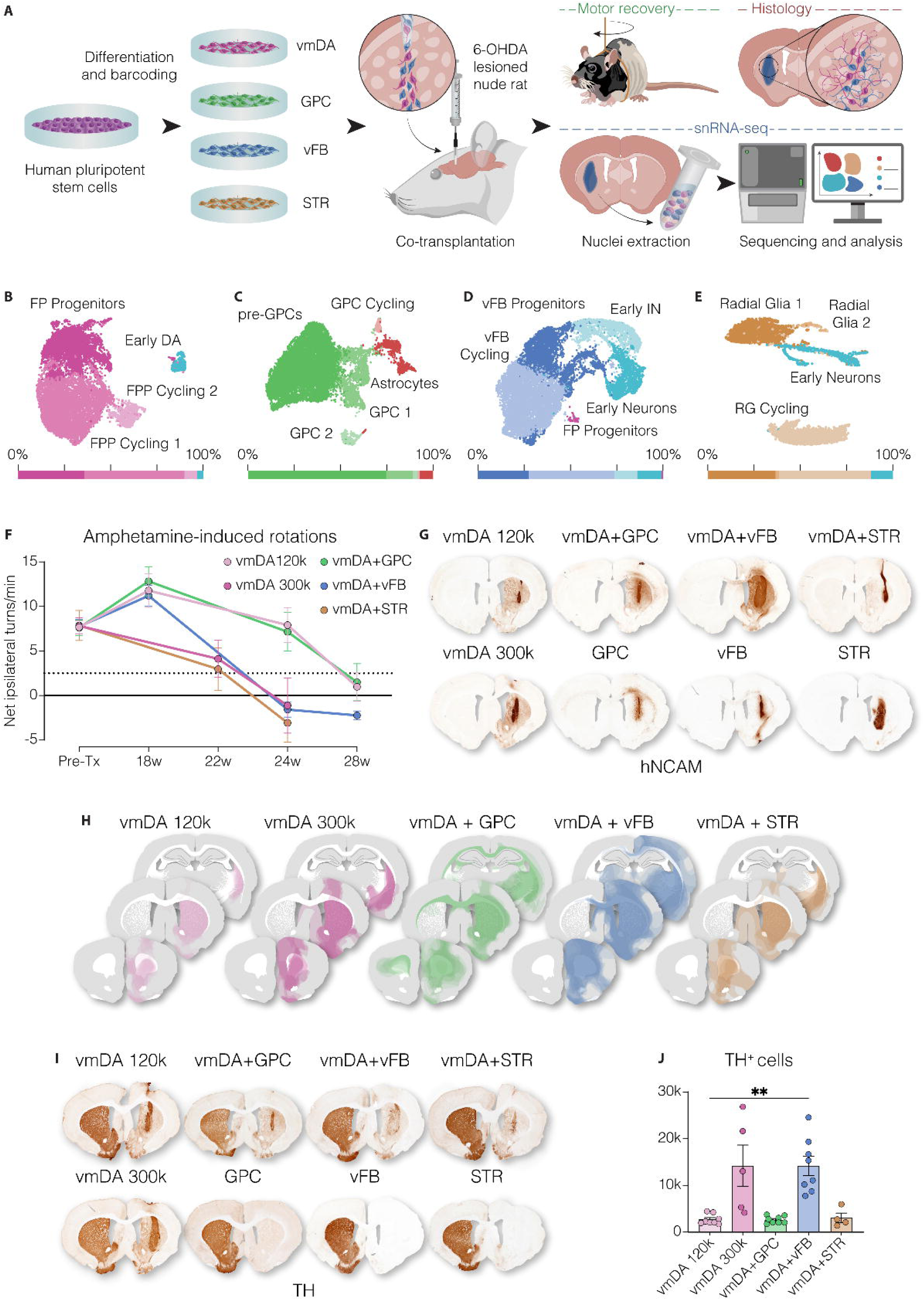
Co-transplantation of vmDA and vFB patterned progenitors leads to an increased yield of TH^+^ neurons. **A)** Schematic overview of co-grafting experimental design. Human pluripotent stem cells were differentiated to vmDA, GPC, vFB and STR progenitors, tagged with inheritable barcoding libraries and co-grafted into the striatum of a preclinical model of Parkinson’s Disease. Post-transplantation analysis included histology, behavioural motor tests and single nuclei RNA sequencing. 6-OHDA, 6-hydroxydopamine. **B-E)** Uniform manifold approximation and projection (UMAP) embeddings of single-cell RNA-seq data from **B)** vmDA (n =10,463 cells), **C)** GPC (n =13,685 cells), **D)** vFB (n =13,858 cells) and **E)** STR (n =5,220 cells) cell preparations prior to transplantation. Clusters are coloured by annotated cell type, and their relative proportions are shown in the bar plots underneath. FPP, floor plate progenitors; DA, dopaminergic neurons; IN, interneurons; RG, radial glia. **F)** Amphetamine-induced rotation test showing motor recovery in all groups receiving vmDA progenitors, with recovery dynamics varying by co-grafted cell type (vmDA 120k n =10; vmDA 300k n = 8; vmDA+GPC n = 10; vmDA+vFB n = 10; vmDA+STR n = 8). At 24 weeks, vmDA+vFB vs vmDA 120k p = 0.0191, vmDA+STR vs vmDA 120k p = 0.0076; ANOVA with Tukey’s multiple comparisons test. **G)** Representative DAB-developed hNCAM immunohistochemistry of coronal striatal sections from all groups (including GPC n = 5; vFB n = 6; STR n = 10). **H)** Composite overlay showing hNCAM^+^ fibers innervating the host brain from all animals in each group. **I)** Representative TH staining in coronal sections of grafted brains. **J)** Stereology quantification of TH^+^ cells 7 months post-transplantation (Kruskal-Wallis with Dunn’s multiple comparisons test, p = 0.0077).

To determine the cell composition, patterning and proliferative state of each cell preparation pre-transplant, we processed them for single-cell RNA sequencing (vmDA, 10,463 cells; GPC, 13,685 cells; vFB, 13,858 cells; STR, 5,220 cells). Transcriptomic profiling confirmed correct regionalization of the progenitors (vmDA, GPC, vFB and STR, respectively), and identified cells in different cycling stages, as well as a small fraction of post-mitotic populations, in all four differentiations (Fig. 1B-E and Suppl. Fig. 1L-S). Cell cycle genes such as *TOP2A*, *MKI67,* and *CENPF* marked cells in active cell division, whose occurrence was comparable in all four batches (62% in vmDA, 63% in GPC, 54% in vFB and 65% in STR, G2M+S) (Suppl. Fig. 1L-O). vmDA progenitors were transplanted into the striatum of 6-OHDA lesioned immune-deficient athymic nude rats, either alone in a low dose (120,000 cells; n = 10) or standard dose (300,000 cells; n = 8), or co-grafted with different supportive progenitor cells. In the co-grafting groups, vmDA progenitors (120,000□cells) were mixed with 180,000□cells of either GPCs (n□=□10), vFB progenitors (n□=□10), or STR progenitor cells (n□=□8), keeping the total number of transplanted cells constant at 300,000 per graft. Control animals were also transplanted with the same preparations of GPC (n = 5), vFB (n = 6) or STR progenitors (n = 10) alone (300,000 cells each). Motor recovery was assessed by amphetamine-induced rotations at 18-, 22-, 24- and 28-weeks post-grafting. Animals transplanted with vmDA progenitors, either alone or co-transplanted with non-DA support cells, exhibited progressive behavioural recovery, while no recovery was observed in animals grafted with support cells alone (Fig. 1F and Suppl. Fig. 2A). We observed that two groups, the vmDA+vFB co-graft and vmDA+STR co-graft mediated an earlier motor recovery compared to animals grafted with the same number of vmDA progenitors alone (120,000 cells), and in the same timeframe as when grafting 300,000 vmDA progenitors (Fig. 1F; at 24 weeks, vmDA+vFB vs vmDA 120k p = 0.0191, vmDA+STR vs vmDA 120k p = 0.0076; ANOVA with Tukey’s multiple comparisons test). Five to eight animals per group were perfused and processed for histology 24-28 weeks after transplantation. Staining for hNCAM confirmed graft survival and maturation in all groups, albeit with differences in graft volume between groups (Fig. 1G and Suppl. Fig. 2B). Mapping the projections from the grafted cells showed extensive innervation to the dorsolateral striatum (dlSTR) in all animals (Fig. 1H and Suppl. Fig. 2C). Human fibers were also observed in other brain areas in a pattern that varied depending on the kind of support cells used for co-grafting (Fig. 1H and Suppl. Fig. 2C). Tyrosine hydroxylase (TH), the rate-limiting enzyme in DA synthesis commonly used to detect DA neurons, was detected only in grafts containing vmDA progenitors (Fig. 1I). This indicates that only these progenitors have the capacity to mature into DA neurons *in vivo*. Quantifications of TH^+^ neurons revealed a ∼5.2-fold increase in the vmDA+vFB co-grafts compared to the control group receiving the same number of vmDA progenitors without vFB cells (Fig. 1J). This increase confirmed the observation from the dose-finding pre-clinical study^5^ and was restricted to the vmDA+vFB co-grafts, suggesting a cell type-specific effect of vFB progenitors on DA neuron formation (Fig. 1J).

### Lineage commitment of vmDA progenitors is affected by co-grafted cells

To characterize the cellular composition of the mature grafts at the molecular level, we performed single-nucleus RNA sequencing (snRNA-seq) on brain tissue dissected from three independently transplanted rats per group, resulting in a total of 56 samples and 116,524 high-quality human nuclei. To ensure consistency with previous studies from our group analyzing vmDA grafts^19–21^, we first examined the vmDA-only grafts. These contained mature DA neurons (positive for *TH*, *SNAP25*, *MYT1L*, *RBFOX3*/NeuN), floor plate progenitors (*NES, CORIN*, *NTN1*), astrocytes (*AQP4, GFAP, GJA1*), GPC (*OLIG1*, *PDGFRA*, *SOX10*) and vascular leptomeningeal cells (VLMCs - *DCN*, *COL1A1*, *COL3A1*), with few or no cycling cells detected in the graft, matching previous observations (Fig. 2A,B and Suppl. Fig. 3A,B). We next analysed the sequencing data from all experimental groups combined (i.e. the co-graft groups plus the individually grafted control groups). This revealed increased diversity of cell types as well as a greater heterogeneity within the neuronal cluster (Fig. 2C,D and Suppl. Fig. 3C). While neurons and astrocytes were present in most grafts (Fig. 2E), VLMCs were detected only in vmDA-containing grafts, suggesting a regional origin of this cell type specifically linked to the vmDA lineage (Fig. 2E and Suppl. Fig. 3C). Joint integration of heterogenous datasets imposes a shared low-dimensional structure, and accordingly, a subset of cells may be pulled toward transcriptionally similar neighbours. Consistent with this, a small number of cells clustered as neurons in the GPC transplant group (Fig 2E). However, in depth analysis of the cells in this cluster confirms that they are immature glial cells and not neurons.

**Figure 2.**
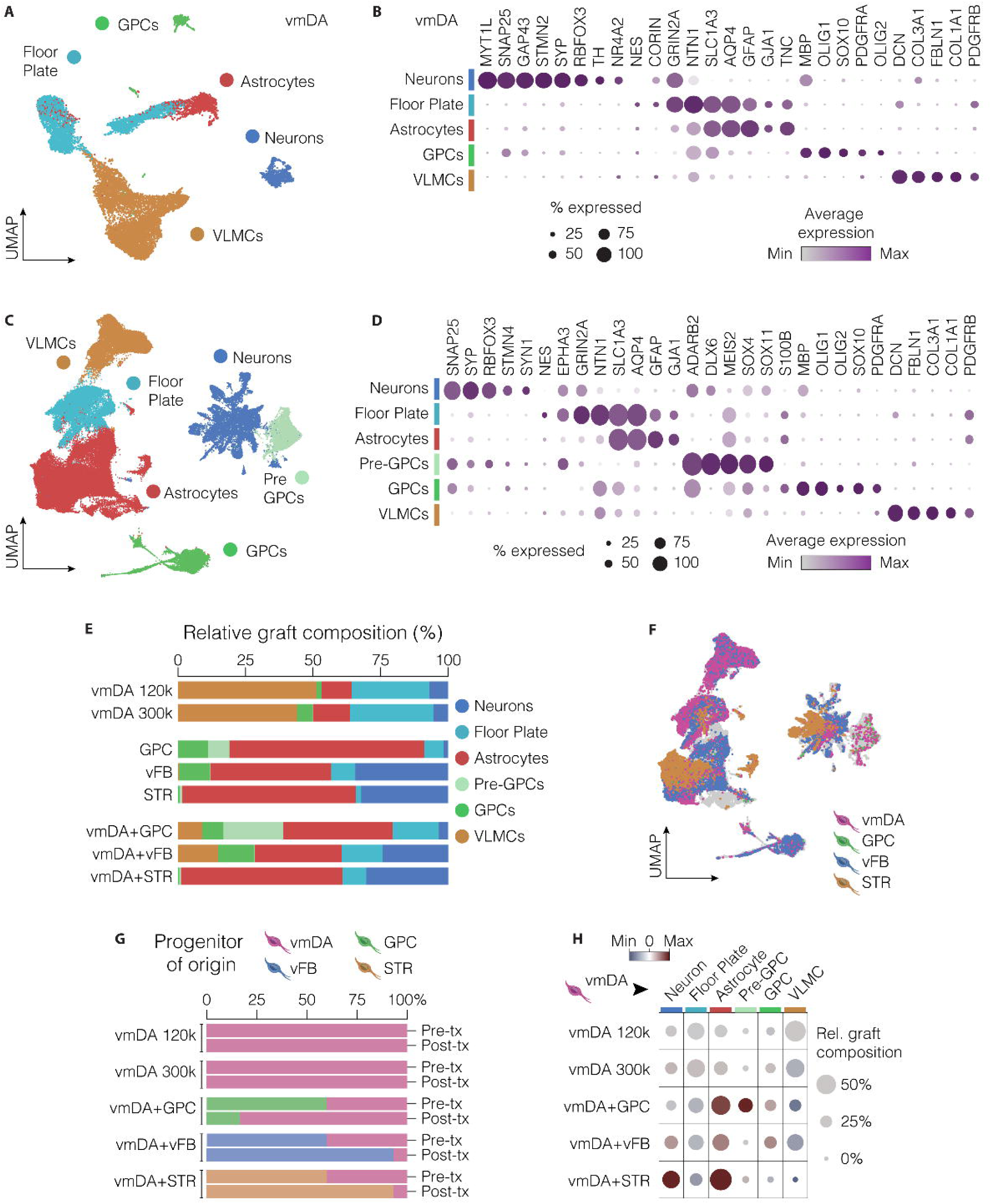
Single-nuclei RNA profiling reveals co-graft cellular composition and modulation of vmDA lineage commitment. **A)** UMAP visualization of vmDA graft composition 28 weeks after transplantation (13,828 cells and 8 samples). GPCs, glial progenitor cells. VLMCs, vascular leptomeningeal cells. **B)** Dot plot showing expression levels of representative marker genes in vmDA graft clusters. **C,D)** Integrated UMAP plot displaying the cellular composition of all grafts 28 weeks after transplantations and dot plot depicting the expression levels of key genes used for cluster annotation (116,524 cells and 56 samples). **E)** Bar plots showing the percentage of nuclei belonging to each cluster across transplantation groups. **F)** UMAP plot showing the progenitor of origin of co-grafted cells 7 months after transplantation, according to the library ID contained in the molecular barcodes used to tag them. Cells lacking a confidently assigned barcode are shown in grey. **G)** Bar plots representing the percentage of human cells originated from the 4 distinct progenitor batches before (top, *Pre-tx*) and after (bottom, *Post-tx*) transplantation. **H)** Dot plot displaying deviations in mature vmDA-derived cell type abundance in the different co-grafts relative to pure vmDA grafts (red = increased relative abundance; blue = reduced relative abundance).

To assign each cell type in the mature graft to its progenitor type of origin, we utilized the distinct sequences contained in the barcode libraries to individually tag the original progenitor populations (Fig. 2F and Suppl. Fig. 3D). Additionally, because vmDA and STR progenitors were derived from different hESC lines (RC17 and H9, respectively), we leveraged genetic demultiplexing to validate their relative abundance in vmDA+STR co-grafts, and overall, our barcode-based methodology (Suppl. Fig. 3E). This analysis revealed substantial differences in the relative contributions of the various progenitors to the mature graft (Fig. 2F,G). STR and vFB-derived cells comprised over 90% of their respective co-grafts, even though the cells were mixed in a 40:60 ratio before grafting (Fig. 2F,G). In contrast, GPC-derived cells in vmDA+GPC co-grafts (mixed at the same 40:60 ratio in the cell preparation used for transplants) represented only 17% of the total cells in the area where the graft was placed, likely due to the migratory nature of GPCs (Fig. 2F,G). Using a barcode-based demultiplexing approach, we further examined how co-grafted cells might influence the differentiation potential of the vmDA progenitors. Differential abundance analysis of mature cell types derived solely from vmDA progenitors showed an increase in neuron and astrocyte populations across all co-grafts, with the most prominent effects observed in the vmDA+STR group (Fig. 2H).

### Neuronal diversity in the co-grafts

To gain a better understanding of the neuronal diversity of the mature grafts originating from each progenitor type, we performed a high-resolution analysis of 23,635 cells within the neuron cluster of the combined dataset. We identified a group of early neurons (*SLC1A3, TNC, SLC4A4*) and 13 postmitotic neuronal clusters (all expressing *MAPT*, *STMN2*, *NCAM1*) (Fig. 3A,B). As previously reported^20,21,24^, DA neurons (*TH*, *LMX1A*, *NR4A2*, *SLC18A2/*VMAT2, *EN1*) were the dominating neuronal subtype derived from vmDA progenitors and were also detected in all co-grafts of vmDA progenitors with supportive cells (Fig. 3B-F). Most of the other neuron subclusters originating from vFB and STR progenitors expressed genes involved in GABA production (*GAD1, GAD2*), in combination with regional markers of the medial (MGE), lateral (LGE) or caudal (CGE) ganglionic eminences (Fig. 3B,D-F). vFB progenitors generated mainly MGE interneurons (*NKX2-1, LHX6, SST, NPY*) at different maturation stages, CGE neurons (*NR2F2,* also known as COUP-TFII*, RARB, PROX1, RORB*) and a small population of cholinergic neurons (*LHX8*, *CHAT*, *NTRK1*, *ISL1* and *SLC5A7* – the choline transporter) (Fig. 3A,B,D-F). On the other hand, GABAergic neurons originating from STR progenitors mostly belonged to the MSN lineage (*BCL11B*, known as CTIP2, *NPAS1*, *SP8*) with a small percentage of striatal interneurons (*ISL1*, *SIX3*, *LAMP5* but *NKX2-1* negative) (Fig. 3A,B,D-F). Furthermore, STR progenitors also matured into a population of glutamatergic neurons (*SLC17A7*, also known as vGLUT1, CTIP2 and *TBR1*) when grafted alone, which was drastically reduced in the co-grafts with vmDA cells (Fig. 3 A,B,D-F).

**Figure 3.**
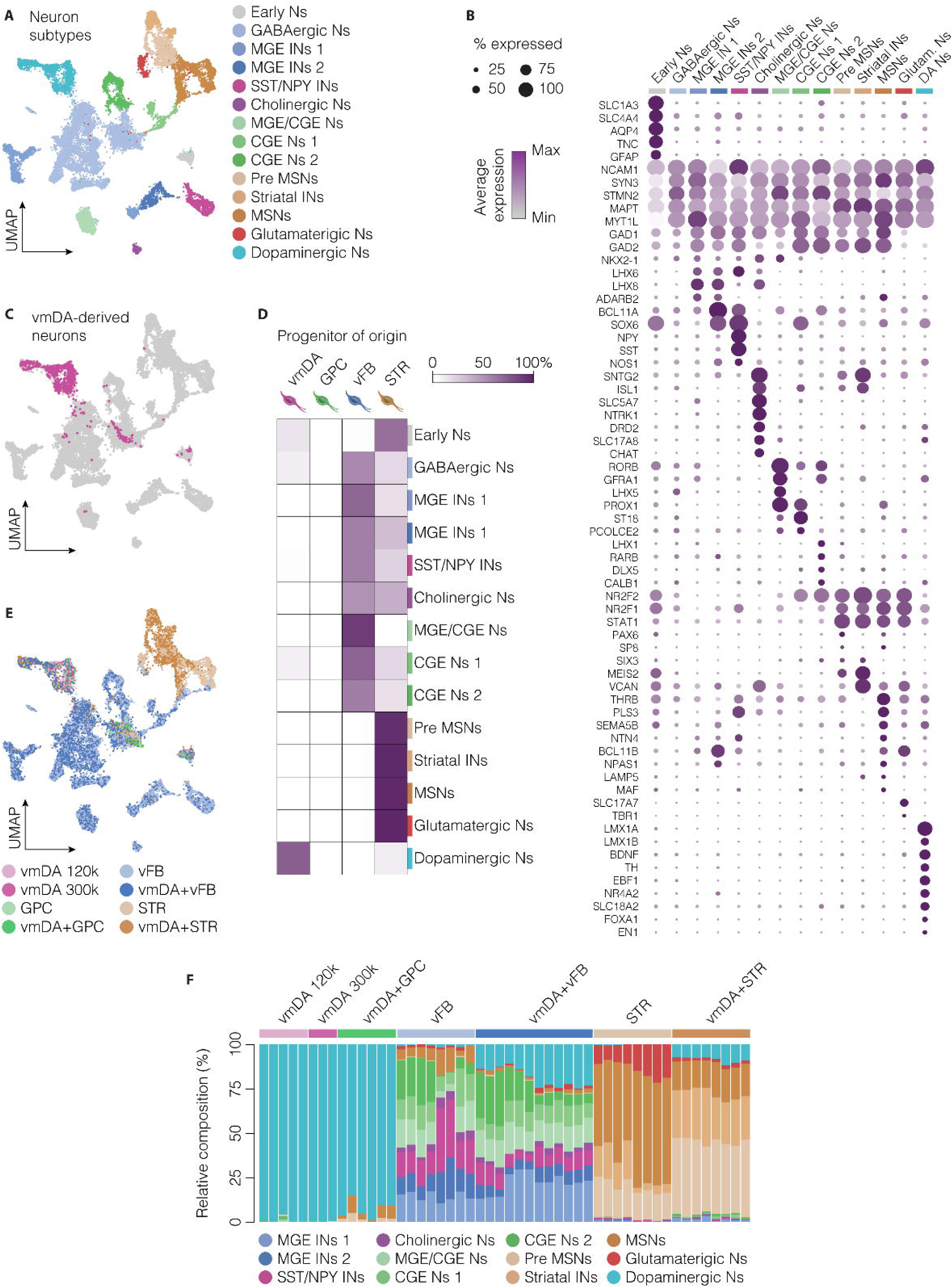
Neuronal diversity in co-grafts. **A)** UMAP visualization of neuronal diversity in co-grafts (23,635 cells). Ns, neurons; MGE, medial ganglionic eminence; IN, interneuron; SST, somatostatin; NPY, neuropeptide Y; CGE, caudal ganglionic eminence; MSN, medium spiny neuron. **B)** Dot plot showing selected markers used for annotating neuronal subtypes. **C)** Visualization of vmDA-derived neurons within the neuronal cluster. **D)** Heatmap depicting the relative proportion of neuronal subtypes originating from each of the four grafted cell preparations. **E)** UMAP embeddings of neuronal clusters, coloured by experimental group. **F)** Bar plot showing the percentage of mature neuronal subtypes composing the neuronal cluster across different samples.

### Increased DA neuron maturation and A9 subtype specification in co-grafts with striatal target cells

Histological analysis confirmed the presence of mature and subtype-specific DA neurons in both vmDA transplants and co-grafts containing vmDA cells, as shown by immunofluorescence of TH in combination with ALDH1A1, GIRK2 and OTX2 (Fig. 4A-C and Suppl. Fig. 4A-C). Quantifications of the co-expression of ALDH1A1, GIRK2 and OTX2 with TH (Fig. 4D-F) revealed that co-transplants with STR progenitors contained higher proportions of ALDH1A1^+^ and GIRK2^+^ cells within the TH^+^ population compared with the other groups (Fig. 4D,E). Conversely, the fraction of OTX2^+^ DA neurons in vmDA+STR co-grafts was significantly reduced compared to vmDA grafts alone (Fig. 4F).

**Figure 4.**
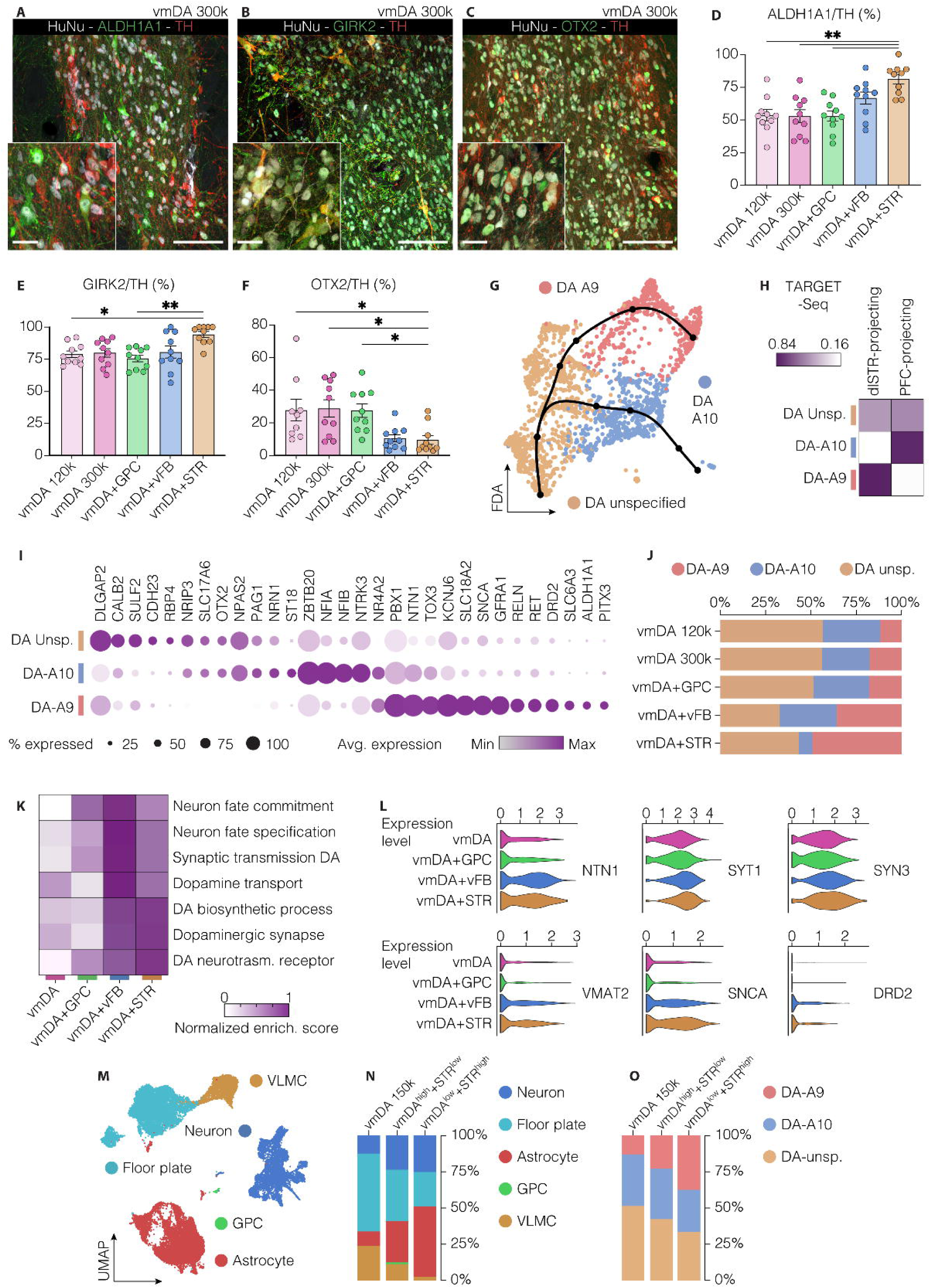
Co-grafting vmDA and STR progenitors influences DA neuron maturation and subtype specification. **A-C)** Immunohistochemistry of ALDH1A1 **A)**, GIRK2 **B)** and OTX2 **C)** coupled with TH and HuNu showing DA neuron subtype diversity in vmDA grafts. Scale bar = 100 μm; inset, 25 μm. **D-F)** Percentage of ALDH1A1- **D),** GIRK2- **E),** OTX2-positive **F)** cells over TH in vmDA-alone grafts and co-grafts. vmDA+STR vs vmDA 120k: ALDH1A1 p = 0.0053, GIRK2 p = 0.0269, OTX2 p = 0.0449, Kruskal-Wallis with Dunn’s multiple comparisons test. **G)** Force-directed k-nearest neighbour graph (SPRING) of computationally sorted DA neurons, with slingshot trajectories (black lines) indicating maturation paths toward A9- and A10-like molecular profiles. **H)** Heatmap showing the correlation between DA neuron subtypes identified in this study and the transcriptomic profiles of dlSTR-projecting or PFC-projecting DA neurons described in Fiorenzano et al. 2024^19^. **I)** Dot plot showing expression levels of selected differentially expressed genes across DA neuron subtypes. **J)** Relative abundance of DA neuron subtypes across co-grafts. **K)** Heatmap showing gene set enrichment analysis (GSEA) scores for gene ontology terms related to DA neuron maturation across experimental groups. **L)** Violin plots depicting expression levels of selected synaptic markers in DA neurons. **M, N)** UMAP visualization and bar plots showing the proportions of distinct cell types across the homotopic transplantation groups. **O)** Bar plots depicting the relative abundance of unspecified, A9-like, and A10-like DA neurons in homotopic grafts at 7 months post-transplantation.

Since single markers are insufficient to reliably distinguish A9 from A10 DA neuron subtypes^31^, we subset the DA neuron cluster from the snRNA-seq dataset and performed a re-clustering, which resolved three transcriptionally distinct DA neuron populations (Fig. 4G). To assign subtype identities to these clusters, we leveraged our recently published TARGET-seq resource^19^, which provides molecular profiles of transplanted human DA neurons with defined projection patterns to either the dlSTR (A9 target) or the prefrontal cortex (PFC, A10 target) (Fig 4H). In particular, the red cluster aligned with the A9/dlSTR-projecting profile, displaying enrichment for *KCNJ6/*GIRK2, *ALDH1A1*, *SLC18A2/*VMAT2, *SNCA*, *SLC6A3*/DAT, *PITX3*, *DRD2*, *GFRA1*. The blue cluster corresponded to A10/PFC-projecting DA neurons, enriched for *OTX2*, *NFIA/B*, *ZBTB20*, *LMO3* (Fig.□4G-I, Suppl.□Fig.□4D–H). The yellow cluster represented an immature DA state characterized by the expression of *CALB2*, *NRIP3*, *SLC17A6/*VGLUT2, and reduced *TH/NR4A2* expression (Fig 4G-I and Suppl Fig. 4D-H), consistent with early developmental trajectories in which DA neurons have not yet acquired target-specific axonal identities^19,32^ (Fig 4H,I).

In vmDA-only and vmDA+GPC co-grafts, most DA neurons retained an immature transcriptional signature at this stage (yellow cluster, Fig. 4G-J and Suppl. Fig. 4I,J). In contrast, major differences were seen when vmDA cells were co-grafted with vFB or STR progenitors. In vmDA+vFB co-transplants, 36% and 32% of the DA neurons were classified as A9- and A10-like, respectively (Fig. 4J and Suppl. Fig. 4I,J). On the other hand, 49% of the DA neurons in the grafts derived from vmDA+STR co-grafts matched the A9/dlSTR-projecting molecular signature, while only 7% belonged to the A10-like cluster (Fig. 4J and Suppl. Fig. 4I,J). This finding suggests that when vmDA progenitors develop alongside their STR targets within the graft, they preferentially mature into A9-like DA neurons, resulting in a greater proportion of cells acquiring the molecular features of the STR-projecting subtype.

Finally, gene set enrichment analysis (Fig. 4K) revealed that DA neurons in vmDA+vFB or vmDA+STR grafts upregulated pathways linked to neuronal fate commitment, DA biosynthesis, and synaptic maturation, with increased expression of functional DA synapse markers such as *SLC18A2/*VMAT2, *DRD2, SNCA*, and *NTN1* (Fig. 4L). These findings support that the co-transplantation with STR target cells not only bias subtype specification toward A9 identity but also promote transcriptional maturation of engrafted DA neurons.

To validate these findings, we conducted a new set of experiments in which vmDA and STR progenitors were mixed in two different ratios - vmDA^low^ + STR^high^ (40:60, same as previous experimental setup) and vmDA^high^ + STR^low^ (80:20) - and co-grafted into the midbrain (homotopic site) of our preclinical PD model. As in the heterotopic intrastriatal grafts, tissue was collected for sequencing 28 weeks post-transplantation. Consistent with previous observations, the homotopic vmDA grafts contained DA neurons, astrocytes, floor plate cells, GPCs and VLMCs (Fig 4M,N). Furthermore, we validated that co-grafting vmDA and STR progenitors promoted DA neuron maturation and A9-like subtype specification, as shown by a dose-dependent decrease in the proportion of not-yet specified DA neurons (yellow cluster) and a parallel increase of A9-like neurons (red cluster) (Figure 4O). Together, our data support the notion that co-grafting vmDA with their STR targets modulates both cellular graft composition and DA neuron subtype specification.

### Transcription factors *EBF3* and *PBX3* orchestrate DA neuron maturation and mature subtype specification

The distinct molecular signatures of A9- and A10-like neurons suggest that human DA neuron specification may be driven by a specific set of transcription factors (TFs). To test this, we constructed gene regulatory networks (GRNs) for DA neurons (Fig. 5A) based on differentially expressed genes that define A9 and A10 neuron clusters (Fig. 5B). This analysis identified key TFs that regulate broad gene modules, including markers associated with A9/dlSTR- and A10/PFC-projecting DA neurons analysed previously (Fig. 5A,B). TFs upregulated in A10-like DA neurons included *NFIB* and *NFIA*, which directly repress the A9-enriched GDNF receptor *GFRA1*, and *TCF4*, a mediator of the Wnt-β-catenin pathway, consistent with previous studies^19,33^ (Fig. 5A,B). Another TF enriched in A10-like neurons was *CREB5,* which inhibits GIRK2/*KCNJ6* expression while activating *ZBTB20*. TFs enriched in A9 DA neurons included *EBF3*, directly controlling *BNC2*, the axon guidance factor *ROBO2* and *SNCA* (Fig. 5A,B). To investigate molecular changes associated with DA neuron maturation, we performed a comparative analysis between unspecified and mature, subtype-specific A9 and A10 DA neurons (Fig. 5C,D). *PBX3* emerged as the central element of this network, regulating the transcription of both *EBF3* and *NFIB*, as well as *ERBB4*, *NTN1* (involved in axon guidance) and *KCNJ6*, *GFRA1* and *FOXP2* (Fig. 5C,D).

**Figure 5.**
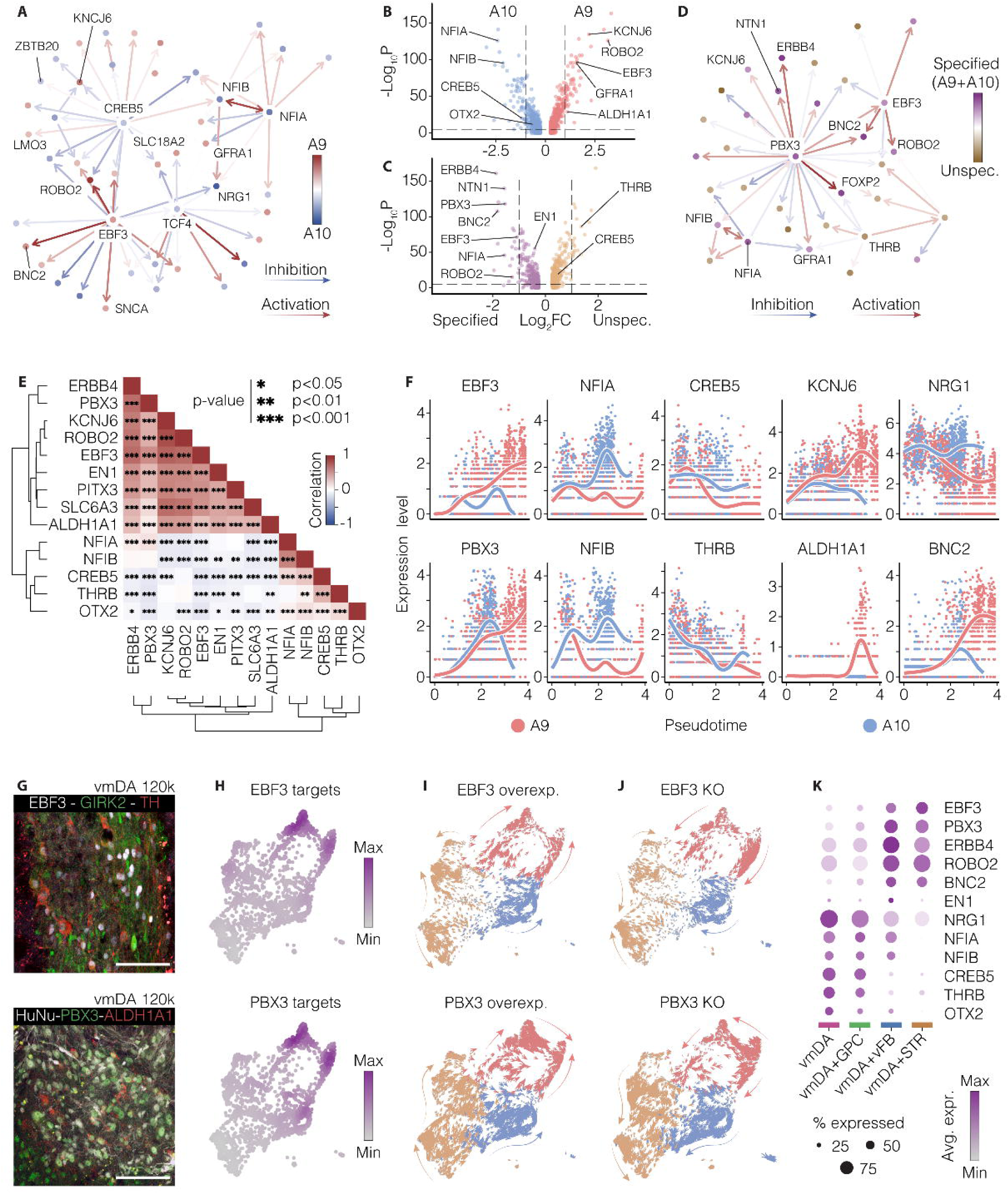
Gene regulatory network analysis identifies transcription factors *EBF3* and *PBX3* as candidate drivers of DA neuron specification and maturation. **A)** Gene regulatory network (GRN) of differentially expressed genes between A9-like and A10-like DA neuron clusters. Nodes represent genes and are colored according to the fold change (Log_2_FC) values shown in the volcano plot in (B). **(B,C)** Volcano plots of differentially expressed genes in A9-like (red) versus A10-like (blue) DA neurons (B), and in specified (A9-like and A10-like together) versus unspecified DA neuron clusters (C). The horizontal dashed line indicates the significance threshold (p < 0.05), and the vertical dashed lines represent the fold-change cutoffs. **D)** GRN of differentially expressed genes between specified (A9-like and A10-like combined) and unspecified DA neuron clusters. Both GRNs (in A, D) were constructed with CellOracle. **E)** Heatmap showing co-expression patterns of selected genes identified in the GRN and mature DA neuron markers. **F)** Expression levels of selected markers along the pseudo-temporal axis showing differential expression along identified A9/A10 trajectories (see Fig. 4G). **G)** Immunohistochemistry of EBF3/GIRK2/TH (top) and HuNu/PBX3/ALDH1A1 (bottom) in vmDA-derived grafts, showing nuclear expression of both EBF3 and PBX3 in human DA neurons. Scale bar = 100 μm. **H)** Feature plot depicting the cumulative expression pattern of *EBF3* (top) and *PBX3* (bottom) targets on the DA neuron SPRING plot. I,J) *In silico* simulation of *EBF3* and *PBX3* overexpression **I)** and knock-out (KO) **J)** obtained with CellOracle. The perturbation effect is shown as a vector overlaid onto the DA neurons SPRING plot and colored based on the subtype assignment. **K)** Dot plot showing the differential expression of selected markers in vmDA-derived DA neurons generated from vmDA grafts alone or in co-grafts with support cells.

The genes identified in the GRNs segregated into two main groups based on their expression patterns. The first group, including *EBF3*, *PBX3*, *ROBO2*, and *ERBB4*, was positively correlated with A9 markers such as *ALDH1A1* and *KCNJ6*, as well as mature DA markers including *SLC6A3*/DAT, *PITX3*, and *EN1* (Fig. 5E). In contrast, the expression of TFs *CREB5* and *THRB* showed an inverse correlation with these markers and was instead associated with A10-enriched genes, including *NFIA*, *NFIB*, and *OTX2* (Fig. 5E). We then examined the temporal expression patterns of the identified TFs and their targets by constructing pseudo time trajectories (Suppl. Fig. 5A) culminating in either A9/dlSTR- or A10/PFC-projecting DA neurons. While most genes showed similar expression levels at the start of the trajectory, differential expression patterns emerged as DA neurons became specified towards A9 or A10 subtypes (Fig. 5F and Suppl. Fig. 5A,B). Importantly, this branching occurred earlier for the TFs *EBF3* and *PBX3* (towards A9 fate) and *NFIA* and *NFIB* (towards A10 fate), preceding the expression of classical markers such as *ALDH1A1* or *KCNJ6* (Fig. 5F and Suppl. Fig. 5B). The centrality of *EBF3* and *PBX3* in the GRN and their early distinct expression profiles suggest a potential role as early transcriptional regulators of mature DA subtype specification.

To expand on our findings, we first confirmed the presence of these TFs in the nuclei of grafted DA neurons (Fig. 5G), as well as in the *substantia nigra* of adult rats (Suppl. Fig. 5C,D). Next, we analyzed the combined expression of all known targets of the two TFs, revealing a pattern specific to dlSTR-projecting DA neurons (Fig. 5H). To further investigate the roles of *EBF3* and *PBX3* in DA neuron specification, we performed *in silico* perturbations of the inferred GRN using CellOracle (Fig. 5I,J). Simulated overexpression of *EBF3* and *PBX3* produced similar results, promoting the specification of mature A9 subtype-specific DA neurons (Fig. 5I). Conversely, *in silico* knockout (KO) simulations of *EBF3* and *PBX*3 increased the proportion of unspecified DA neurons lacking target-specific innervation^19^, with similar dynamics observed for known DA TFs *EN1* and *OTX2* (Fig. 5J and Suppl. Fig. 5E-H).

The identification of TFs involved in DA neuron maturation and specification prompted us to investigate whether these factors might be upregulated through interactions between different cell types within the co-grafts. To explore this, we compared the transcriptomes of vmDA progenitor-derived DA neurons co-grafted with either GPCs, vFB, or STR progenitors. Interestingly, A9/dlSTR-associated TFs and their downstream targets, including *EBF3, PBX3, ERBB4*, *BNC2* and *ROBO2*, were upregulated in DA neurons maturing in co-grafts with vFB and STR cells (Fig. 5K and Suppl. Fig. 5I,J). Conversely, A10 TFs such as *NFIA, NFIB*, and *OTX2* were downregulated in DA neurons co-grafted with STR progenitors (Fig. 5K and Suppl. Fig. 5I). Additionally, TFs associated with more immature states, like *CREB5* and *THRB*, were preferentially expressed in DA neurons grafted alone or with GPCs, with minimal transcriptional differences between these two groups (Fig. 5K and Suppl. Fig. 5K).

### Transcription factors *EBF3* and *PBX3* promote DA neuron maturation *in vitro*

We next tested whether forced expression of the candidate TFs *EBF3* and *PBX3* could promote the maturation of VM neurons *in vitro*. First, we confirmed that both TFs are expressed in DA neurons within VM organoids (Fig. 6A,B) and then induced their overexpression using lentiviral open reading frames (ORFs) starting at day 60 of differentiation, at a stage when DA neurons are already generated but have not yet acquired a mature subtype identity^34^. Organoids overexpressing either *PBX3* or *EBF3* exhibited a transcriptional shift toward a more mature DA neuron phenotype, characterized by increased expression of mature DA markers and reduced expression of early developmental genes compared to controls (Fig. 6C and Suppl. Fig. 6A,B).

**Figure 6.**
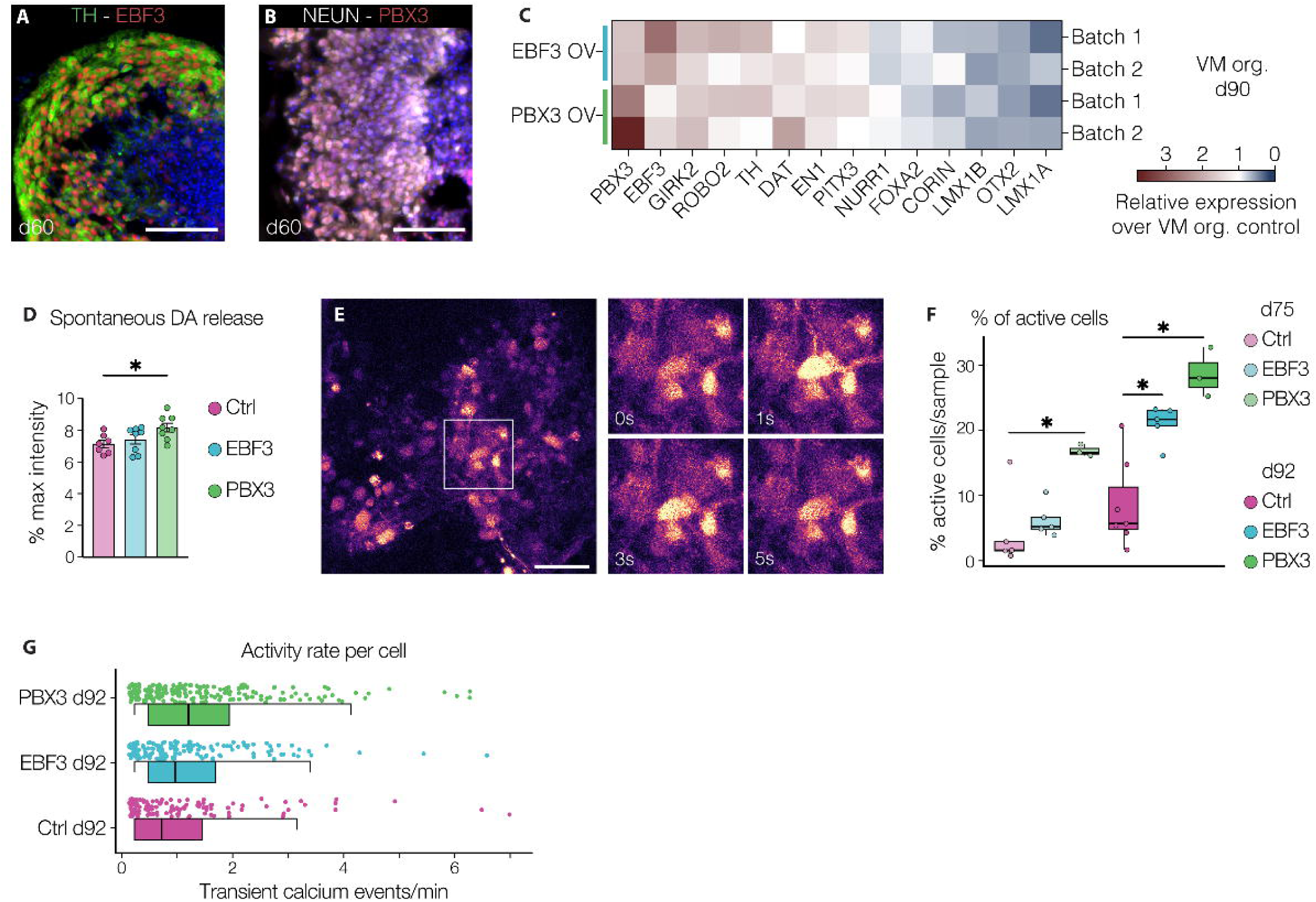
*EBF3* and *PBX3* promote DA neuron maturation and functional activity in VM organoids. **A,B)** Immunohistochemistry of **A)** TH/EBF3 and **B)** PBX3/NeuN in VM organoids at day 60 of differentiation. Scale bars = 100 µm. Nuclei are stained with DAPI. **C)** Expression levels of key marker genes involved in DA neuron maturation and specification in two independent batches of VM organoids overexpressing *EBF3* or *PBX3* at day 90 (n = 4; 3 individual organoids per sample). Expression values are reported as fold change relative to untransduced controls from the same batch. **D)** Quantification of DA in day 90 VM organoids transduced with *EBF3* or *PBX3* overexpression vectors and corresponding controls (n = 8; p = 0.0355; one-way ANOVA with Tukey’s test). **E)** Representative fluorescence-intensity image of intracellular Ca²□levels in a day 90 VM organoid, with four snapshots illustrating temporal changes in intracellular fluorescence. Scale bar = 50 µm. **F)** Box plot showing the percentage of active cells - defined as cells exhibiting at least one transient calcium peak during a 5-minute recording - per organoid at day 75 and day 90. In total, 28 organoids and 5,942 ROIs were processed (Ctrl d75 n = 5; Ctrl d92 n = 7; EBF3 n = 5; PBX3 n = 3; d92 PBX3 OV vs Ctrl p = 0.017; d92 EBF3 OV vs Ctrl p = 0.012; d75 PBX3 OV vs Ctrl p = 0.036; Wilcoxon rank-sum test). **G)** Combined box–scatter plot reporting the activity rate of active cells per minute (n = 127 cells for Ctrl; n = 152 for EBF3; n = 201 for PBX3; PBX3 vs Ctrl p = 0.00024; Wilcoxon rank-sum test).

To determine whether these transcriptional changes translated into functional effects, we performed live calcium imaging and DA release assays using DA sniffer-cells^35,36^ to monitor spontaneous DA release. We observed that *PBX3*-overexpressing (PBX3-OV) organoids release significantly more DA than controls, providing direct evidence that *PBX3* enhances presynaptic DA functionality (p = 0.036, one-way ANOVA with Tukey’s test; Fig. 6D). Calcium imaging further revealed that both PBX3-OV and EBF3-OV organoids presented an increased fraction of active cells, with PBX3-OV exerting a stronger effect (day 75, PBX3 p = 0.036; day 92, EBF3 p = 0.012; day 92, PBX3 p = 0.017; Wilcoxon rank-sum test; Fig. 6E,F). Moreover, cells in PBX3-OV organoids exhibited a higher frequency of calcium events per minute compared to controls, indicating increased spontaneous activity consistent with advanced maturation (Fig. 6G). Together, these findings demonstrate that elevating *EBF3* or *PBX3* expression in developing VM organoids accelerates both molecular and functional maturation of DA neurons, with *PBX3* producing the strongest effects on spontaneous neuronal activity and DA release.

## Discussion

Stem cell-based replacement therapy represents a promising strategy to restore DA transmission in the striatum by replacing the DA neurons lost in PD, and these efforts have now advanced to clinical trials^10,12,13^. Refined cell differentiations and improvements of intrinsic donor cell qualities as well as manufacturing strategies are continuously being developed^10,37–40^. In parallel, it is increasingly recognized that understanding host-graft interactions and how cellular composition guides the differentiation, maturation and function of transplanted DA neurons can also lead to grafts with optimized therapeutic effects.

Along these lines, a number of studies with rodent fetal cells have shown that co-grafting fetal VM tissue with cells with neurotrophic properties, including astrocytes and fetal striatal tissue, promotes the survival of the transplanted DA cells and, in some cases, enhances innervation and more robust functional recovery in transplantation models of PD (reviewed in Rodríguez-Pallares et al., 2023)^41^. Although much fewer studies have been conducted using stem-cell derived DA neurons, co-grafting with astrocytes has shown to increase DA neuron yield and improves therapeutic outcomes in preclinical models of PD, due to the neurotrophic properties of the co-grafted astrocytes^42^. Another study suggests that co-transplanting DA progenitors with astrocytes may be used as a strategy to limit potential spreading of α-synuclein from the host into the transplants^43^. Of great interest is also a recent study showing that co-grafting DA progenitors with T cells (Treg) dampens the inflammatory response associated with the grafting procedure, leading to increased graft survival. This effect has been attributed, at least in part, to the neuroprotective effects that Treg cells exert on the grafted cells^44^. Together, these studies support the notion that co-grafting additional cell populations can modulate vmDA graft outcomes and underscore the increasing focus on refining graft composition and transplantation paradigms to enhance clinical efficacy.

To explore additional strategies to improve the therapeutic benefits of DA grafts, we investigated how the cellular microenvironment within the graft and specific co-grafted cell types modulates lineage commitment, functional maturation and subtype specification of transplanted DA neurons. To address this, we co-transplanted vmDA progenitors with GPCs, vFB or STR neurons into a well-established 6-OHDA preclinical xenograft model, revealing cell-type specific effects effects on the DA neurons within the grafts. In particular, we found that animals grafted with vmDA progenitors mixed with either vFB or STR cells showed earlier motor recovery compared to animals grafted with the same number of vmDA progenitors alone, an effect not observed in co-grafts with GPCs. Interestingly, quantifications of DA neurons 6 months post-grafting revealed that co-grafts containing vFB cells had significantly more DA neurons, while co-grafts containing STR did not, suggesting distinct mechanisms underlying faster motor recovery. In vFB co-grafts, the earlier recovery likely reflects increased DA neuron survival and DA neuron content, whereas in STR co-grafts, it likely results from enhanced maturation and/or altered DA subtype specification.

To investigate the molecular basis of these effects, we performed snRNA-seq on mature grafts coupled with molecular barcode-tagging to identify the progenitor of origin of each mature cell type in the resulting grafts^20^. This revealed that DA neurons in the co-transplants exhibit a higher expression of genes associated with neuron fate commitment and specification, as well as gene ontology (GO) terms specifically linked to DA neuron maturation, compared to DA neurons grafted alone or with GPCs. The transcriptomic analysis also revealed that although co-transplants with vFB interneurons increased the total number of mature subtype-specific DA neurons, the relative proportions of A9-like and A10-like DA neurons remained unchanged at approximately 50:50. In contrast, co-grafting with STR cells specifically increased the number of A9 neurons and decreased the number of A10 neurons, suggesting that target acquisition with STR cells promotes A9 specification of the DA neurons within the vmDA co-grafts. These findings align with previous work showing that the absence of striatal target cells in the host brain significantly reduces the number of A9-like DA neurons in DA grafts, while increasing the number of A10-like neurons^45,46^. Together, these data support the idea that contact with their appropriate developmental targets refines the molecular profile of mature DA neurons, promoting their specification towards the A9 and A10 subtypes^19,31,46^.

Our molecular approach enabled an in-depth analysis of the genes and GRNs of the grafted DA neurons, allowing us to identify regulators of DA neuron development. This included genes like *ERBB4*, expressed in the mouse SNpc independently of *Nurr1* and *En1*, and *NTN1*, a guidance cue involved in segregating SNpc- and VTA-derived axons^47–49^. Furthermore, A9-like enriched genes included *ROBO2,* associated with axon guidance of SNpc DA neurons; *SNCA*, encoding α-synuclein and implicated in PD; and the GDNF receptor *GFRA1*, which promotes striatal innervation from A9 DA neurons^33,50^. In contrast, PFC-projecting DA neurons were enriched for the Wnt-pathway mediator *TCF4* and the TF *NFIB*, aligning with previous studies^19^. Among the identified genes, the TFs *EBF3* and *PBX3* emerged as potential central regulators of DA neuron maturation and specification based on their early and subtype-enriched expression. Previous studies in mice established a critical role for *Ebf3* in midbrain DA neuron development, where gain- and loss-of-function experiments in ES-derived DA neurons showed that Ebf3 selectively promotes DA differentiation and maturation, upstream of key DA determinants such as *Nurr1* and *Pitx3*, while having minimal effects on pan-neuronal genes (*Syp*, *Mapt*). Notably, *EBF3* and a broader network of DA development and maturation – including members of the *PBX* family – are downregulated in SNpc from PD patients, suggesting that the disruption of this regulatory axis may contribute to DA vulnerability^51,52^. Together with our *in silico* perturbation analyses and correlations with classical A9-enriched markers, these findings support a model in which *EBF3* and *PBX3* act as positive regulators of A9 DA neuron maturation. We validated this model in VM organoids, where overexpression of *EBF3* and *PBX3* drove DA neurons toward a more mature molecular and functional identity, characterized by upregulation of neurogenic programs and enhanced neuronal activity, including elevated calcium dynamics and DA release - particularly pronounced in PBX3-overexpressing organoids.

The data presented here provides the first evidence that co-grafting stem cell-derived DA progenitors with defined additional cell types directly affects lineage commitment and molecular specification, ultimately influencing graft composition and DA neuron functionality. A deeper understanding of these cellular interactions within the graft opens new opportunities for therapeutic strategies that target not only the DA neurons themselves, but also the non-DA components of the graft. Together, these insights underscore the value of designing next-generation cell products that leverage cooperative interactions between multiple progenitor populations in rationally designed grafts withs optimized therapeutic effects for maximized clinical efficacy.

## Material and methods

### Human pluripotent stem cell differentiation

GMP-grade RC17 human embryonic stem cells (hESCs) (Roslin Cells, cat. no. hPSC reg RCe021-A), were cultured in iPS Brew XF medium (Miltenyi Biotec) on laminin-521 (0.5 μg/cm² in DPBS with Ca^2+^/Mg^2+^, Biolamina)-coated plates (Sarstedt). Cultured cells were passaged at 70-90% confluency with EDTA (0.5 mM, 7 min at 37 °C) and reseeded at a density of 5,000-10,000 cells/cm² in iPS Brew XF supplemented with ROCK inhibitor (10 μM, Y-27632, Miltenyi, cat. no. 130-106-538) for the first 24 hours after plating. Differentiation toward vmDA and vFB progenitors involved dual SMAD inhibition (10 µM SB431542, Miltenyi, cat. no. 130-106-543; 100 ng/mL Noggin, Miltenyi, cat. no. 130-103-456) for neuroectoderm induction, and activation of SHH pathway (300 ng/mL SHH-C24II, Miltenyi, cat. no. 130-095-727) for ventralization, following the detailed protocol outlined in Nolbrant et al. 2017^28^. In addition, vmDA progenitors were exposed to GSK3 inhibitor (0.9 µM CHIR99021, Miltenyi, cat. no. 130-106-539) and FGF8b (100 ng/mL, Miltenyi, cat. no. 130-095-740) for caudalization. GPCs were also derived from RC17 following the Wang et al. 2013^30^ protocol with the modifications outlined in Nolbrant et al. 2020^53^. STR progenitors were derived from H9 hESC line (WiCell, cat. no. hPSCreg WAe009-A) according to Delli Carri et al. 2013^29^. Five days before transplantations, all four cell preparations were genetically tagged with lentiviral barcode libraries carrying a specific ID, to discriminate between the different cell preparations in co-grafts. Each library contained approximately 10^6^ unique barcodes and transductions were performed at multiplicity of infection (MOI) 1. Detailed generation of the barcode libraries and validation of library diversity are described in Storm et al. 2024^20^. Cells were transplanted after 16 days of differentiation for vmDA and vFB progenitors, 20 days for STR progenitors and 170 days for GPCs. Before surgery, cells were washed in DPBS, centrifuged at 400 g for 5 min, and resuspended in Hanks Balanced Salt Solution (HBSS, Thermo Fisher Scientific, cat. no. 14175095) supplemented with 0.05% DNase (Sigma-Aldrich, cat. no. DN25). All cell preparations used for transplantations met pre-established quality control criteria outlined in the appropriate protocol (Suppl. Fig. 1). To assess GPC differentiation prior to transplantation, cultured cells were analyzed by fluorescence-activated cell sorting (FACS Aria III cell sorter, BD Biosciences, nozzle 100 µm), utilizing fluorochrome-conjugated antibodies against human CD140a (PE anti-human CD140a, BD Biosciences, cat. no. 556002, 1:10) and CD44 (APC anti-CD44, Miltenyi, cat. no. 130-095-177, 1:500). Compensation was performed using single-stained controls, and gating was based on Fluorescence Minus One (FMO) controls.

### Surgical procedures and cell transplantation

All surgical procedures were performed on female athymic nude rats (Hsd:RH-Foxn1^rnu^) under general anesthesia by intraperitoneal (i.p.) injection of a mixture of ketaminol® (ketamine hydrochloride, 45 mg/kg) and domitor® (medetomidine, 0.3 mg/kg) according to weight. Lesion of the nigrostriatal pathway was induced by unilateral injection of 6-hydroxydopamine into the right medial forebrain bundle (MFB), using a total volume of 3 μL at a freebase concentration of 3.5 μg/μL and an infusion rate of 0.3 μL/min. The injection was administered at the following stereotaxic coordinates relative to bregma: anterioposterior (A/P) -3.9 mm, mediolateral (M/L) -1.2 mm, and dorsoventral (D/V) -7.3 mm from dura, with the head positioned flat (dorsoventral difference between bregma and lambda ≤ 0.2 mm). Lesion severity and motor recovery were assessed by amphetamine-induced rotations (i.p. injection of 3.5 mg/kg of amphetamine; Apoteksbolaget, Sweden) at least 4 weeks post-lesion or at 18-week, 22-week, 24-week and 28-week post-transplantation, respectively. Rotational behaviour was recorded for 90 min using an automated rotometer system (RotoMax, Omnitech Electronics), and results were expressed as net turns per minute. For intrastriatal cell transplantations, animals received a total of 300,000 progenitors unilaterally via 2 tracts with 2 deposits per tract (1 μL/deposit, 75,000 cells/ μL, infusion rate 1μL/min, 2 min diffusion time per deposit) at the following coordinates relative to bregma: A/P +1.4 mm, M/L -2.6 mm, D/V (from dura) -4.0/-5.0 mm, and A/P +0.9 mm, M/L -3.0 mm, D/V (from dura) -4.0/-5.0 mm, adjusted to flat skull position. For co-grafts, cell suspensions consisted of 40% vmDA progenitors and 60% support cells (STR, GPCs or vFB progenitors) for a total of 300,000 cells/rat. For homotopic transplantations, animals received a total of 150,000 progenitors in a single tract with 2 deposits at the following coordinates: A/P -4.9 mm, M/L -2.3 mm, D/V (from dura) -5.5/-6.0 mm (flat-head position). Homotopic co-grafts were prepared by mixing 40% vmDA progenitors with 60% STR cells (vmDA^low^+STR^high^) or 80% vmDA progenitors with 20% STR cells (vmDA^high^+STR^low^). At experimental endpoints, animals were deeply anesthetized with sodium pentobarbital (300 mg/kg, Apoteksbolaget, Sweden) and transcardially perfused with 0.9% saline solution followed by ice-cold 4% paraformaldehyde (PFA). All experimental procedures were performed under the EU directive 2010/63 and the Swedish animal protection legislation and were approved by Lund Animal Ethics board and the Swedish Department of Agriculture (Jordbruksverket).

### Nuclei extraction and isolation

Nuclei isolation from long-term grafts was performed as described in Sozzi et al., 2025^54^. Briefly, rat brains were dissected in ice-cold artificial cerebrospinal fluid (aCSF) containing 119 mM NaCl, 26 mM NaHCO_3_, 2.5 mM KCl, 1.25 mM NaH_2_PO_4_, 2.5 mM CaCl_2_, 1.3 mM MgSO_4_, and 25 mM glucose. Coronal sections (375 μm thick; amplitude 1.7 mm, speed 0.20 mm/sec) were cut utilizing a vibratome (Leica VT1200S), and graft-containing regions were isolated under a dissection microscope. Dissected grafts were then transferred to a pre-chilled Dounce homogenizer containing 1□mL of lysis buffer (0.32 M sucrose, 5 mM CaCl_2_, 3 mM MgAc, 0.1 mM Na_2_EDTA, 10 mM Tris-HCl, pH 8.0, 1 mM DTT, 0.1% Triton X) and gently homogenized on ice (10 loose- and 10 tight-fitting pestle strokes). The nuclei suspension was centrifugated at 900 g for 15 min at 4 °C, and the pellet resuspended in 200-400 μL of ice-cold wash buffer (0.1% BSA fraction V, and 0.04 U/µL and 0.02 U/µL RNase inhibitors in PBS). Nuclei were purified by FACS (FACSAria III cell sorter, BD Biosciences, nozzle 100 µm) based on DRAQ7 staining (1:1000) and event size. After sorting, 7,000-1,0000 nuclei per sample were collected in BSA pre-coated DNA LoBind tubes (Eppendorf) for subsequent snRNA-seq library generation.

### Sample preparation for single-cell RNAseq

Progenitors cultured in monolayers were dissociated using Accutase (StemPro; Thermo Fisher Scientific, cat. no. A1110501) for 5 min at 37 °C. After incubation, cells were gently detached and collected in DMEM/F12 medium (Thermo Fisher Scientific, cat. no. 31330095) supplemented with 5% KnockOut Replacement Serum. The cell suspension was centrifuged at 400 g for 5 min, and the resulting pellet was resuspended at a concentration of 1,000 cells/μL in BSA 0.1% (Fraction V) in HBSS (Thermo Fisher Scientific, cat. no. 14175095). A total of 7,000–1,0000 viable cells per sample were subsequently processed for scRNAseq library preparation (n = 2 samples per cell preparation).

### Single cell/nucleus RNAseq (library generation)

Single cell RNA sequencing was used to profile the cellular heterogeneity of progenitors pre-transplantation, while single nuclei transcriptomic was used to assess the graft and organoid composition. Single cells and nuclei were captured using the Chromium platform (10X Genomics, PN-120233) according to the manufacturer’s recommendations and encapsulated with barcoded beads. Cell/nuclei lysis and reverse transcription were performed in a droplet reaction with poly(dT) primers containing cell-specific barcodes, unique molecular identifiers (UMI) and sequencing adaptor sequences. After pooling and amplification, the library was fragmented and processed according to the manufacturer’s protocol. For graft samples, half of the unfragmented library was instead used for double-barcode amplification, as described in Storm et al. 2024^20^, introducing the TruSeq Read 2 sequence, an i7 index and the P7 sequence. After quality control using a Bioanalyzer (DNA, HS kit, Agilent), library preparations were sequenced with Illumina NovaSeq 6000 or Illumina NovaSeq X 2 x 100 bp reads for transcriptome libraries and 2 x 50 bp reads for barcodes libraries.

### Single cell/nucleus RNAseq (alignment, processing and data analysis)

Cell Ranger (version 7.0, 10x Genomics) software was used for base-call demultiplexing and raw data processing. Following demultiplexing of the sequencing data with bcl2fastq (v2.19), reads containing the specific barcode library motif were identified and extracted from the fastq files using a custom Perl script. In the graft dataset presented here, the barcode capture efficiency was of 31.15% (28,598 barcoded cells analysed). For the capture and extraction of the barcodes, please refer to the detail and code published in Storm et al., 2024^20^. For graft samples, fastq snRNA-seq reads were aligned to a combined human/rat reference genome (both version 93 from Ensembl) containing both human (GRCh38) and rat (Rnor 6.0) sequences using STAR aligner as implemented in the cellranger pipeline. The specie of origin was assigned based on the GEM classification analysis from the cellranger output. Preprocessing and downstream analysis were performed with Seurat (v.4.3) and R (v4.2.2) on macOS Ventura v13.2.1. Nuclei with fewer than 700 or more than 10,000 detected genes, as well as those with a mitochondrial fraction exceeding 1%, were excluded from further analysis. After log-transformation, 6,000 highly variable genes were identified with *vst* from Seurat integrated function followed by principal component analysis (PCA, n = 150). To integrate individual 10X runs, the harmony package was applied (by sample, v1.03, lambda = 0.99) and corrected coordinates were used for downstream UMAP projection and clustering. Clusters were identified using the Louvain algorithm, (resolution 0.03, Seurat) and annotated based on conventional markers as well as differentially expressed genes identified by FindAllMarkers function (Seurat). For label transfer from the homotopic transplantation dataset presented in Fiorenzano et al. 2024^19^, anchors and prediction score were identified with FindTransferAnchors and TransferData (Seurat). Genetic demultiplexing of vmDA+STR co-grafts was performed with demuxlet^55^ which utilizes inter-individual genetic variation to assign each cell to its most likely donor in pooled experiments. To generate reference genotype data, we prepared whole-genome sequencing libraries from individual cell lines using the Illumina DNA PCR-Free Prep kit. These libraries were then sequenced on an Illumina NovaSeq X platform, achieving an average coverage depth of 30x. Bioinformatic analysis of the resulting whole-genome sequencing data was conducted using the Sarek pipeline^56^ and demuxlet (version 1.0.5) on the BAM files generated by cellranger. The resulting hPSC line classifications were further analyzed downstream in R. Neuron and DA neuron clusters subsetting were followed by an additional harmony integration and reclustering (resolution 0.28 and 0.07, respectively). K-nearest-neighbor graph of DA neurons was rendered using SPRING force-directed layout on normalized expression counts^57^. The Slingshot package (v2.6) was used to perform pseudo-time and trajectory analyses, while the escape function enrichIt (v1.8) was used for gene set enrichment analysis (GSEA). Plots were generated using the ggplot2 package (v3.4.4) or Seurat functions when appropriate. *In silico* perturbation and GRN analysis were conducted using the CellOracle package^27^ (v0.12.1) in Python (v3.8.16) with the base GRN published by Paul et al. 2015^58^. Progenitor fastq reads were aligned exclusively to the human genome (GRCh38) following a similar downstream pipeline. Quality control parameters were uniformed across the 4 datasets, retaining cells with 700-12,000 genes detected and less than 10% mitochondrial reads. Optimal clustering resolution was evaluated using the silhouette function from the cluster package (v2.1).

### Histology and immunohistochemistry

After perfusion, brains were dissected out and post-fixed overnight in 4% PFA at 4 °C, then transferred to 25% sucrose solution for 2-3 days until the brains had sunk to the bottom of the vial. The brains were cut on a freezing microtome into 8 series of 35 μm-thick coronal sections. For DAB-developed immunohistochemistry and immunofluorescence, free-floating sections were incubated overnight at 4 °C in a blocking solution (0.25% Triton-X, 5% serum for the species specific to the secondary antibody in 0.02 M KPBS) containing the appropriate primary antibodies. When required, antigen retrieval was performed with a Tris-EDTA buffer (pH 9.0, 10 mM Tris Base, 1 mM EDTA) for 15 min at 80 °C. Sections were then incubated with secondary antibodies (fluorophore- or biotin-conjugated for DAB detection) for 1 hour in blocking solution and mounted onto gelatin-coated microscope slides. For DAB staining, sections were dehydrated through an ascending ethanol percentage, cleared with xylene and coverslipped using DPX mounting medium. Fluorescent sections were coverslipped using polyvinyl alcohol mounting medium containing DABCO (Sigma-Aldrich). All sections were left to dry overnight before imaging. A list of primary antibodies and dilutions used in this study is reported in Table 1.

**Table 1.**
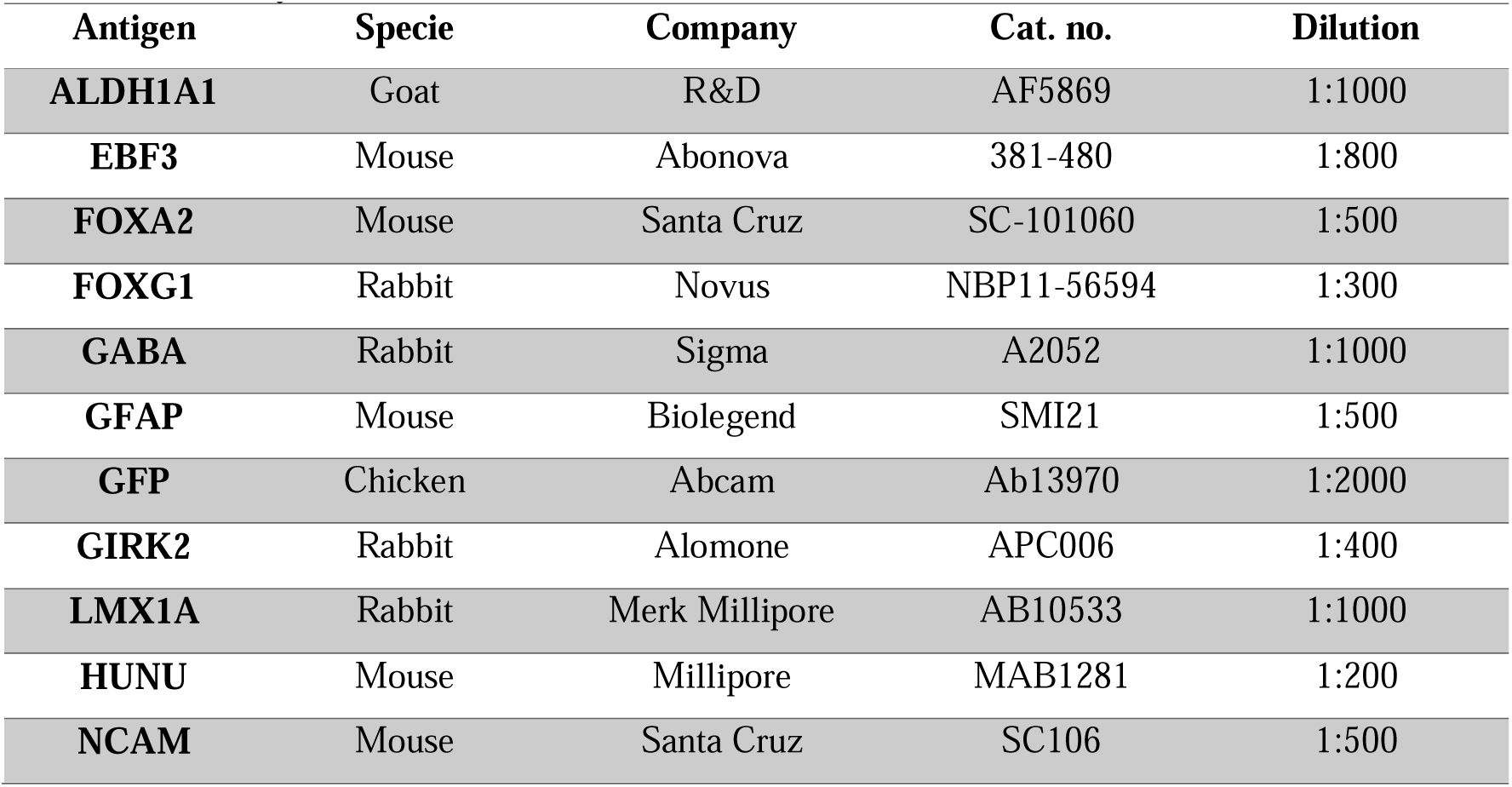
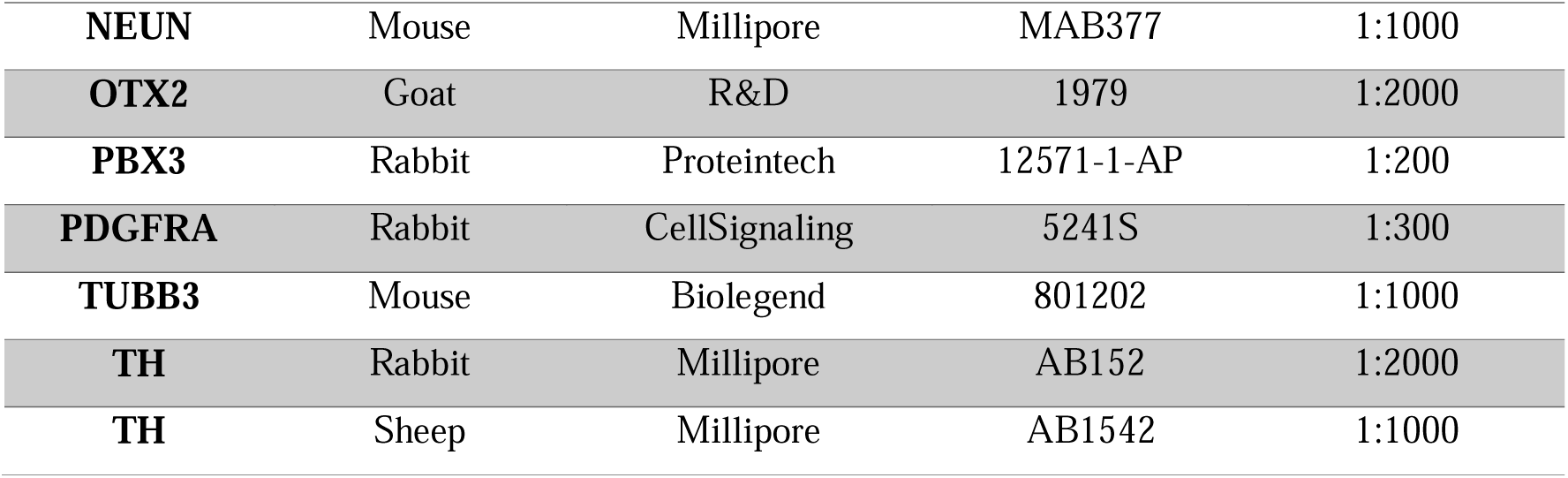
Primary antibodies.

### RNA extraction and qPCR

To analyze gene expression, cell cultures were disrupted using 350 μL of RLT lysis buffer, and total RNA was isolated using the RNeasy Microkit (Qiagen) according to the manufacturer’s protocol. RNA was transcribed into cDNA using the Maxima First Strand cDNA Synthesis kit for real-time quantitative PCR (qRT-PCR) (Thermo Fischer Scientific). cDNA (1 μL, 10 ng/μL) was then mixed with the relevant primers (4 μL, 0.95 μM; Table 2, Integrated DNA Technologies) and with LightCycler 480 SYBR Green Master (5 μL, Roche) in 384-wells plates using a Bravo Automated Liquid Handling Platform (Agilent). mRNA expression levels were determined by qRT-PCR in a LightCycler 480 II instrument (Roche) using a 40-cycle two-step protocol consisting of a 30s denaturation step at 95°C and a 1 min annealing/elongation step at 60°C. Relative gene expression was calculated from three technical replicates using the ΔΔCT method. The average fold-change was calculated using *ACTB* and *GAPDH* as housekeeping genes and the gene expression levels were normalized to undifferentiated RC17 hESC values.

**Table 2.**
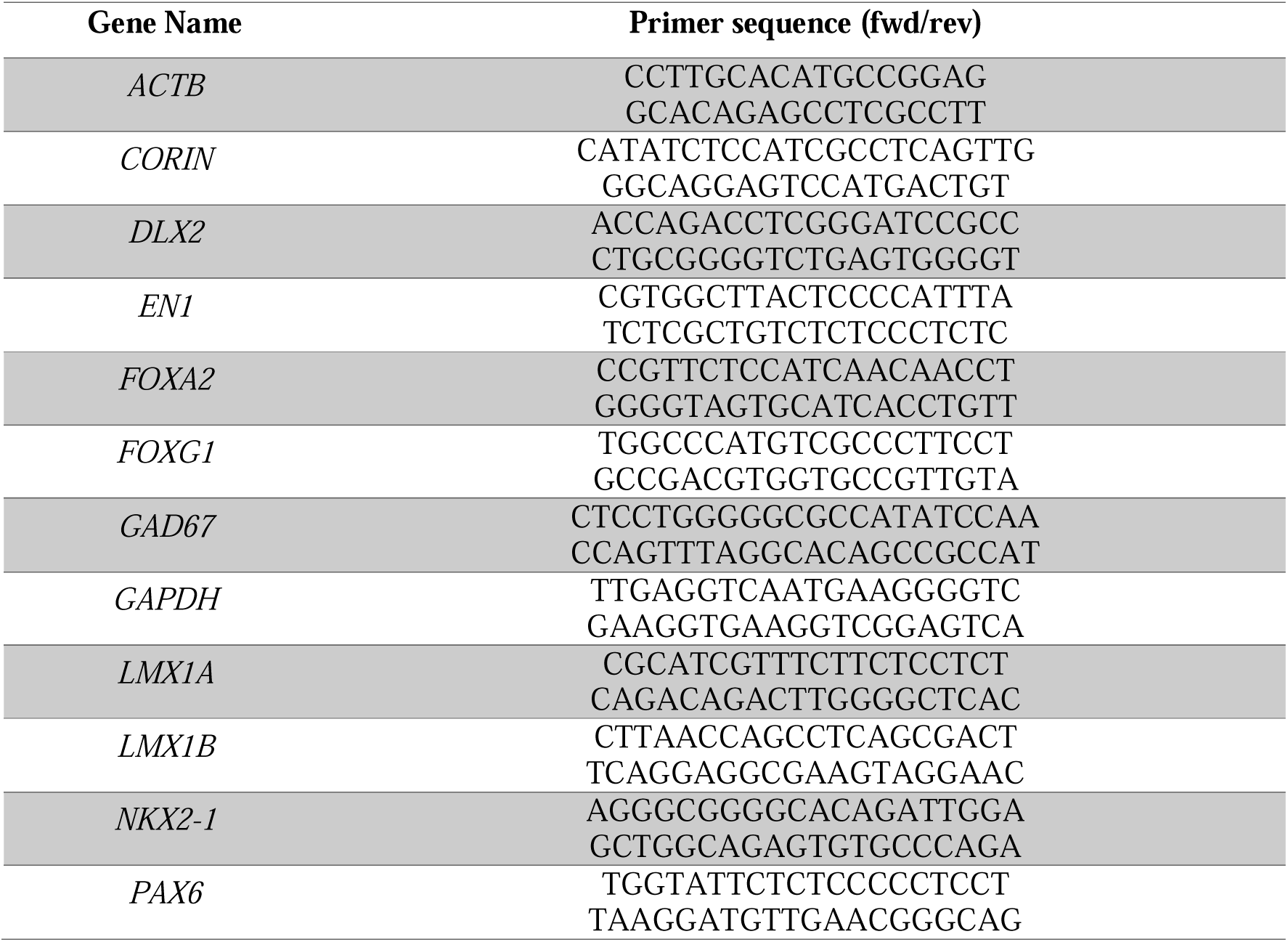
Sequence of qPCR primers.

### Image acquisition and quantifications

Images of hNCAM DAB-developed coronal sections were captured using an Epson Perfection V850 PRO scanner or an Olympus AX microscope. Immunofluorescence pictures of progenitor cells were acquired with a Leica DMI6000B widefield microscope (Leica LAS X software). Graft volume was determined by measuring the area of the graft core in every eighth section throughout the graft using ImageJ software (version 2.9.0). These measurements were then multiplied by the distance between sampled sections and calibrated against a reference scale. Quantification of TH^+^ neurons was conducted in every section with a visible graft under an Olympus microscope (BX53) equipped with an XYZ motorized stage using the Optical Fractionator probe of Stereo Investigator software (version 2023.2.3, MBF Bioscience). Each graft was outlined as a region of interest (ROI) with the 4x objective. Cell profiles were observed with a 100x oil-immersion objective using a 100 μm x 100 μm counting frame. Section thickness was measured every tenth counting frame. The total number of TH^+^ neurons per graft was estimated by the Stereo Investigator Software using the optical fractionator equation^59^. For DA neuron subtype analysis, images were acquired on a Leica TCS SP8 laser-scanning confocal microscope (Leica LAS X acquisition software) using a 20x objective, with a resolution of 2048 x 2048 px, a scan speed of 200 Hz and 1 µm z-step, spanning the full thickness of the 35 μm-thick brain sections. To ensure a comprehensive representation of each graft, three pictures were acquired from the central section of each graft, along with one image from the immediately adjacent rostral and caudal sections to it. Downstream image processing was performed using the ZEISS arivis Pro (v4.2) software, where the volume of HuNu^+^ nuclei was measured in a reconstructed three-dimensional space. Human nuclei with a volume larger than 80 μm^3^ were counted and used to evaluate the concurrent presence of TH^+^- and ALDH1A1^+^/GIRK2^+^/OTX2^+^-cells.

### Ventral midbrain (VM) organoid differentiation

VM-patterned organoids were generated following the protocol described by Sozzi et al., 2022^60^. Briefly, cell aggregates were formed by seeding 8,000 hESCs per well in U-bottom 96-well plates (Corning, cat. no. 7007) using iPS-Brew medium supplemented with 10 μM Y-27632. The differentiation medium consisted of a 1:1 mixture of DMEM/F12 and Neurobasal, supplemented with N2 (1:100), SB431542 (10 μM), rhNoggin (100 ng/mL), SHH-C24II (400 ng/mL), and CHIR99021 (1.5 μM), along with glutamine (2 mM), MEM-NEAA, penicillin-streptomycin (20 U/mL), and 2-mercaptoethanol (50 μM). From day 8 onward, the aggregates were exposed to FGF-8b (100 ng/mL). At day 12, developing VM organoids were transferred to Neurobasal medium supplemented with B27 (–vitamin A, 1:50), BDNF (20 ng/mL), and L-ascorbic acid (200 μM). On day 16, VM organoids were embedded in 30 μL Matrigel droplets and maintained in terminal differentiation medium containing Neurobasal with B27 (–vitamin A), BDNF (20 ng/mL), L-ascorbic acid (200 μM), db-cAMP (500 μM), and DAPT (1 μM) for long-term culture. For validation of *EBF3* and *PBX3* roles in DA neuron development, VM organoids were transduced at day 60 with lentiviral vectors carrying specific ORFs for *EBF3* and *PBX3* at MOI of 2. The MOI was estimated based on live cell counts obtained after dissociation of three individual organoids from the same developmental stage and batch.

### Calcium imaging

Calcium imaging was performed on VM organoids at days 75 and 92 of differentiation. Organoids were incubated with 5 μM Calbryte 520AM (AAT Bioquest cat. no. 20650) in BrainPhys imaging-optimized medium (STEMCELL technologies, cat. no. 05796) supplemented with 0.04% Pluronic F-127 (AAT Bioquest cat. no. 20053) for at least 1 hour at 37°C. Following incubation, organoids were transferred to glass-bottom plates (Ibidi, cat. no. 80807) and imaged using a Leica Stellaris 5 confocal laser-scanning inverted microscope. Time-lapse recordings were acquired at 4 Hz for 5 min using a 40x oil-immersion objective. Individual ROIs were identified in ImageJ (NIH) and raw data was processed and visualized in R (v.4.2.2).

### Dopamine detection

Dopamine detection was performed using HEK-293 Flp-In T-Rex cells stably expressing the fluorescent G protein–coupled receptor-based dopamine sensor GRAB_DA2M_, as previously described^35,36,61^. Cells were maintained in DMEM supplemented with 10% FBS, 15 µg/mL blasticidin, and 200 µg/mL hygromycin B. Forty-eight hours prior to measurements, sniffer cells were seeded in poly-L-ornithine–coated imaging chambers (Ibidi, 18-well) and sensor expression was induced with 1 µg/mL tetracycline. Samples were collected from individual VM organoids incubated for 1 hour in 200 µL of HBSS, snap-frozen, and stored at −80 °C until analysis. Baseline, sample-induced fluorescence response, and the saturated sensor response to 10 µM DA solution was recorded on a Leica widefield microscope using a 20x objective. Image sequences were aligned and mean fluorescence intensity per image quantified using ImageJ (NIH). Results were plotted as %max in Prism (GraphPad).

### Statistical analysis

The normality of data distributions was verified with a Shapiro-Wilk test, and parametric or non-parametric tests were performed accordingly, as indicated in the text or figure legend. Normally distributed data were analyzed with ANOVA with Tukey’s multiple comparisons test or using the Kruskal-Wallis test followed by Dunn’s multiple comparison test otherwise. Outliers were identified using ROUT (Q = 1%) and removed from the dataset. All data are expressed as mean ± standard error of the mean (SEM). Statistical analyses were performed using GraphPad Prism (v. 10.10). For all figures: * p<0.05, ** p<0.01, *** p<0.001.

## Data and code availability

Single-cell/nucleus RNA sequencing data generated in this study are publicly available at the Gene Expression Omnibus (GEO) under accession number GSE312551. The analysis code related to this work is available from the leading author upon request.

## Acknowledgments

We thank Bengt Mattsson, Malin Åkerblom, Ulla Jarl, Jenny G Johansson, Sol da Rocha Baez, Anna Hammarberg, Marie Persson Vejgården and Michael Sparrenius for excellent technical assistance, and Fredrik Nilsson, Jessica Giacomoni, Kerstin Laurin for valuable input. This work was supported by funding to M.P. from HORIZON-ERC-2024-SYG, project 101167102 Custom-Made, Swedish Research Council (VR 2021-00661), Swedish Parkinson Foundation (Parkinsonfonden), Swedish Brain Foundation, the Royal Physiographic Society in Lund (44435 and 43374), The Crafoord Foundation (20250837), and the Strategic Research Areas at Lund University MultiPark (Multidisciplinary research in Parkinson’s disease) and StemTherapy/Lund Stem Cell Center. The authors would also like to acknowledge Clinical Genomics Lund, SciLifeLab and Center for Translational Genomics (CTG), Lund University, for providing sequencing service

## Author contributions

Conceptualization: MP. Methodology: ES, MGG, JM, GRP, ABr, SC, GG, MH, LS, DB, JK, AF, PS. Investigation: ES, MGG, JM, GRP, ABr, SC, GG, MH, LS, DB, AF. Formal Analysis: ES, PS. Visualization: ES. Funding acquisition: MP, PS, ES. Supervision: MP, EC, ABj. Writing – Original Draft: ES, MGG, PS, MP. Writing – Review & Editing: All authors.

---

**Supplementary Figure 1.**
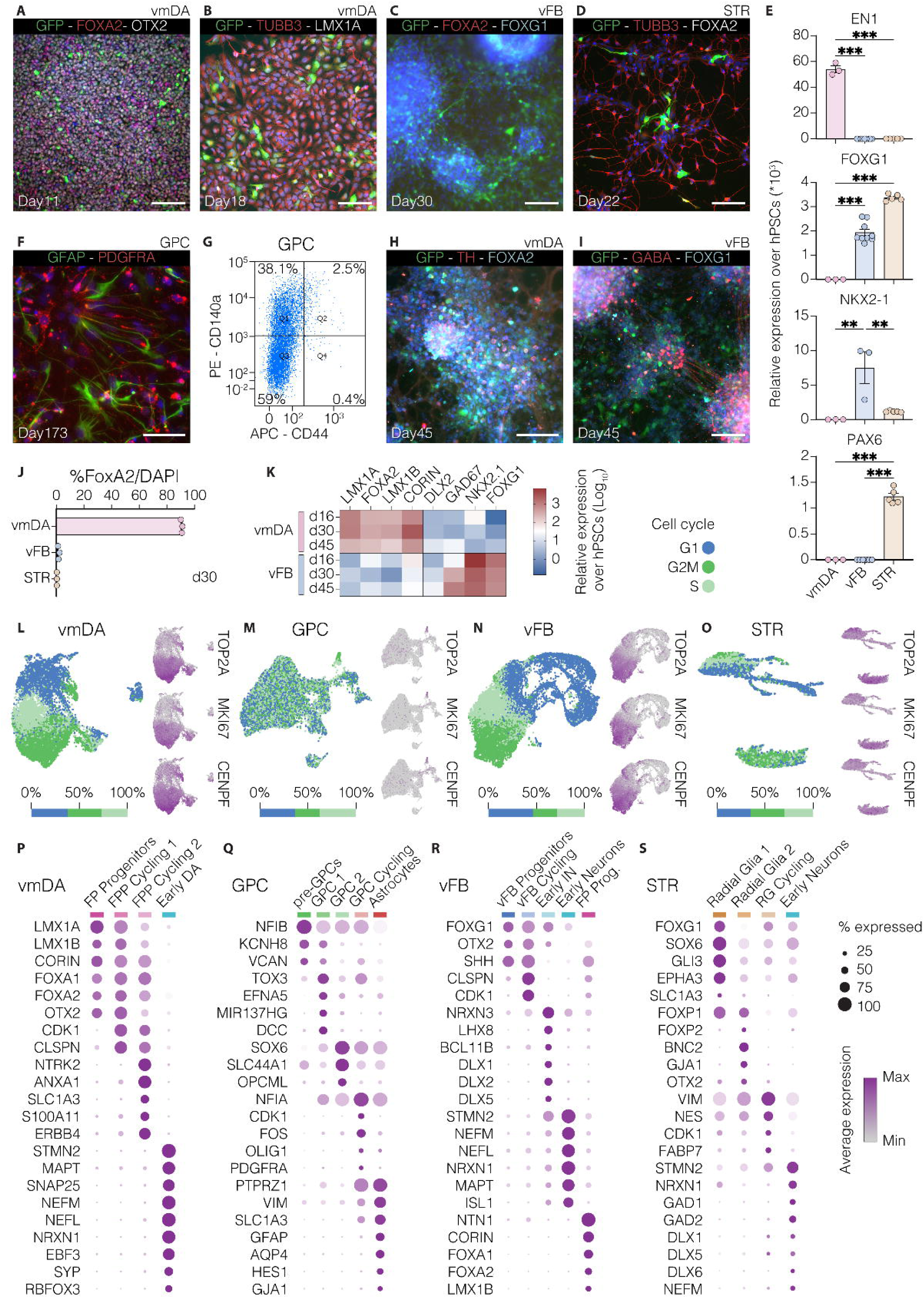
Characterization of vmDA, vFB, GPC and STR progenitors prior to transplantation. Immunocytochemistry for **A)** GFP (barcode)/FOXA2/OTX2 in vmDA progenitors at day 11 and **B)** GFP/TUBB3/LMX1A at day 18 of differentiation. Scale bars = A, 200 μm; B, 100 μm. Immunocytochemistry for **C)** GFP/FOXA2/FOXG1 in vFB progenitors at day 30 and **D)** GFP/TUBB3/FOXA2 in STR progenitors at day 22 of differentiation. Scale bars = 100 μm. **E)** Relative qRT-PCR expression level of regional markers in vmDA (pink), vFB (blue) and STR (yellow) cell preparations at the time of transplantation, normalized to hPSC values. **F)** GFAP/PDGFRA staining of GPC cell preparation. Scale bar = 100 μm. **G)** Representative FACS plot showing CD140a and CD44 expression in GPCs before transplantation. Immunocytochemistry for **H)** GFP/TH/FOXA2 in vmDA cultures and for **I)** GFP/GABA/FOXG1in vFB cultures at day 45 of differentiation. **J)** Quantification of FOXA2^+^ cells at day 30 in vmDA, vFB and STR-patterned progenitors (n = 3). **K)** Relative gene expression (qRT-PCR) of vmDA and vFB markers at multiple differentiation stages (day16, day30, day45). **L-O)** Predicted cell cycle phase distribution, percentage of cycling cells and feature plots of cell cycle markers across the 4 cell preparations: vmDA **L);** GPC **M);** vFB **N);** STR **O).** G1, blue; G2M, green; S, light green. **P-S)** Dot plot illustrating differentially expressed genes and manually selected markers in vmDA **P)**, GPC **Q)**, vFB **R)**, and STR **S)** patterned progenitors.

**Supplementary Figure 2.**
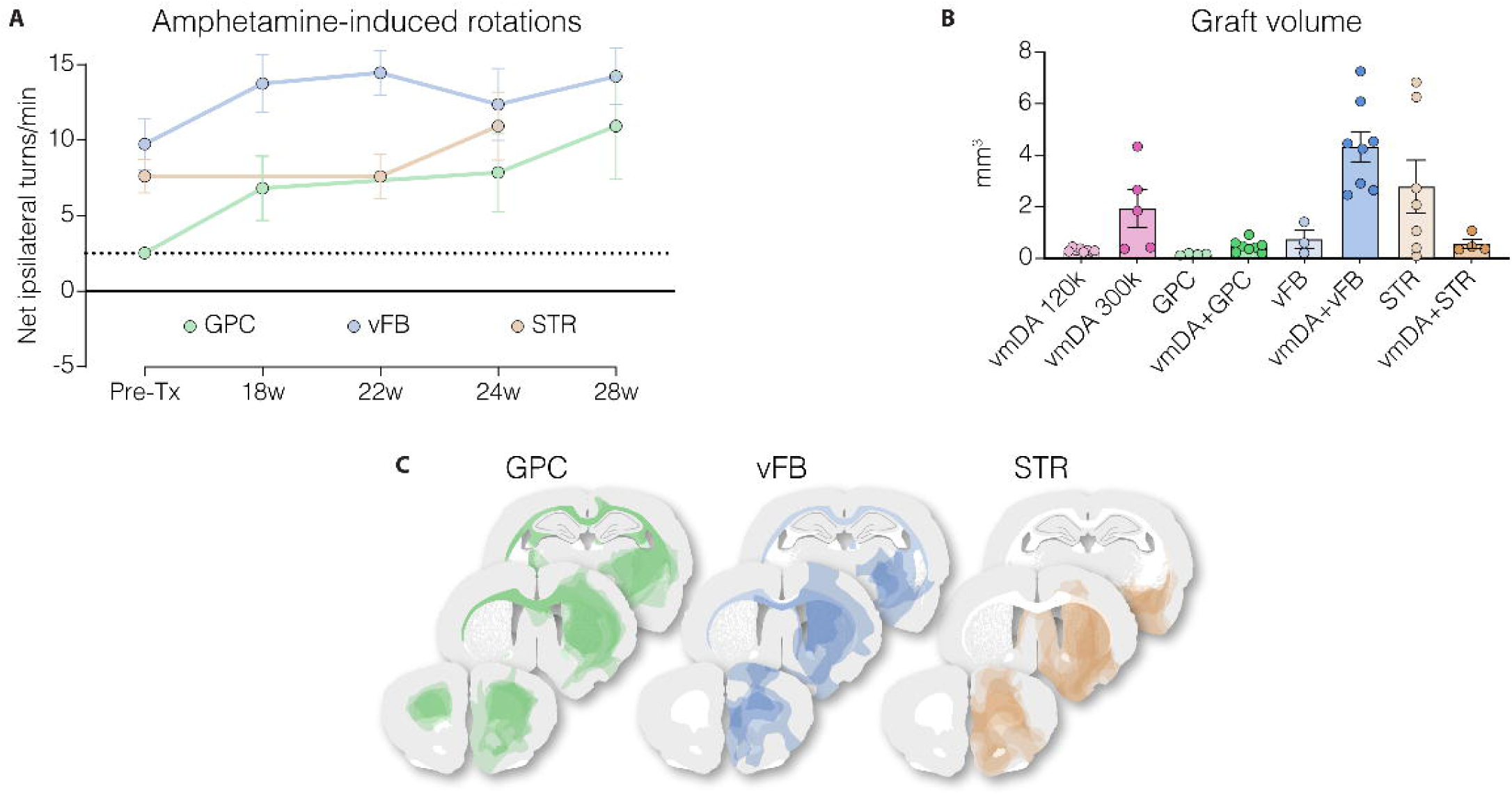
Support-cell-only grafts do not induce motor recovery. **A)** Amphetamine-induced rotation test indicating lack of motor recovery in animals grafted with support cells only. **B)** Quantification of graft volume across all experimental groups; each dot represents one animal. **C)** Composite overlay of hNCAM^+^ areas from all animals grafted with control cells alone.

**Supplementary Figure 3.**
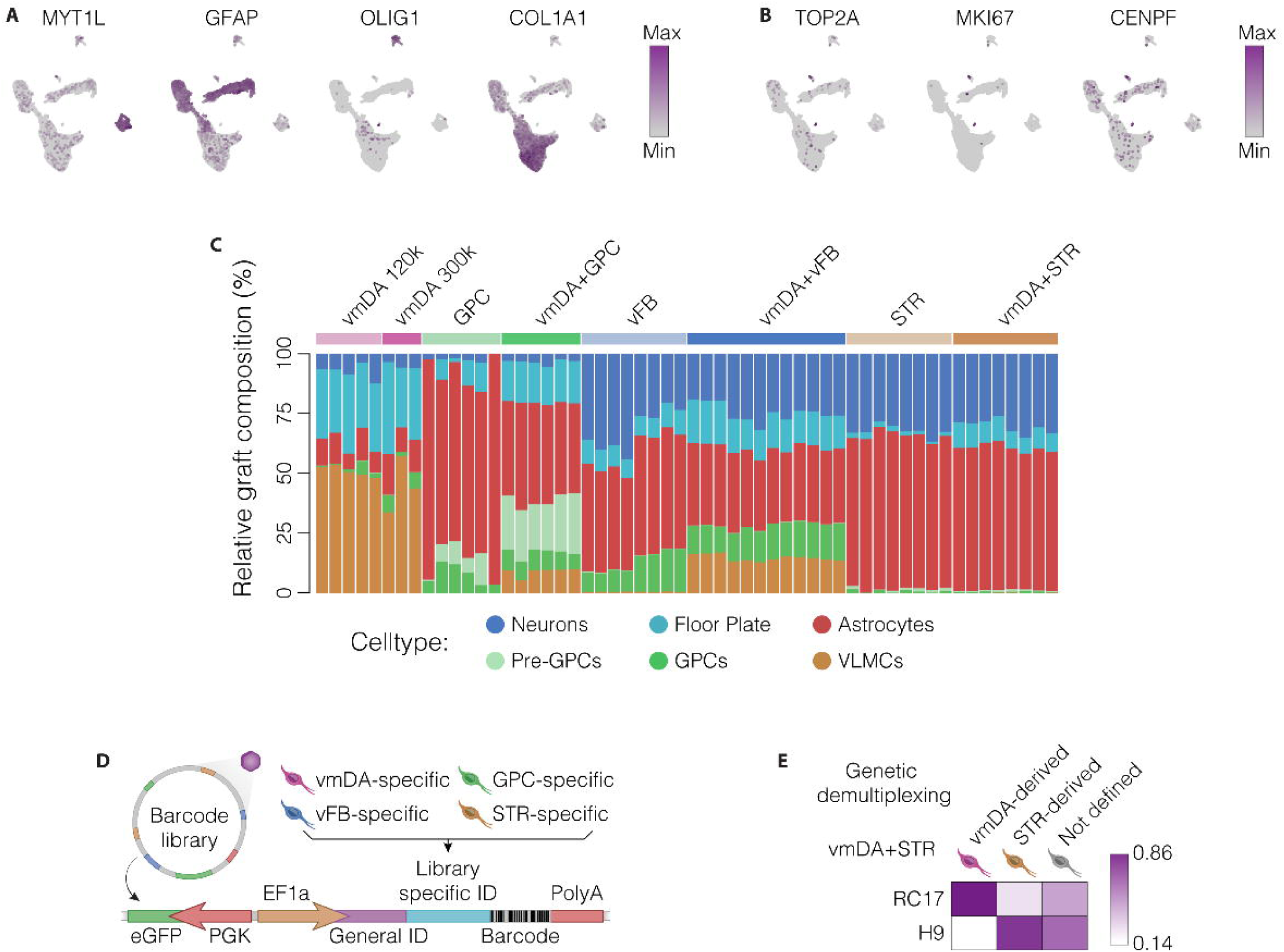
Cellular composition and heterogeneity of vmDA grafts. **A)** Feature plots of key markers for neurons, astrocytes, GPCs and VLMCs and **B)** cycling cells in vmDA grafts. **C)** Bar plots depicting the relative abundance of each cluster across all experimental groups, separated by sample. **D)** Schematic of the barcode library design used to identify distinct progenitor populations; eGFP is driven by the PGK promoter, and the progenitor-specific barcode ID is driven by the EF1a promoter. **E)** Heatmap showing the correlation between the barcode-based demultiplexing of vmDA+STR co-grafts and the genetic demultiplexing based on the cell line of origin.

**Supplementary Figure 4.**
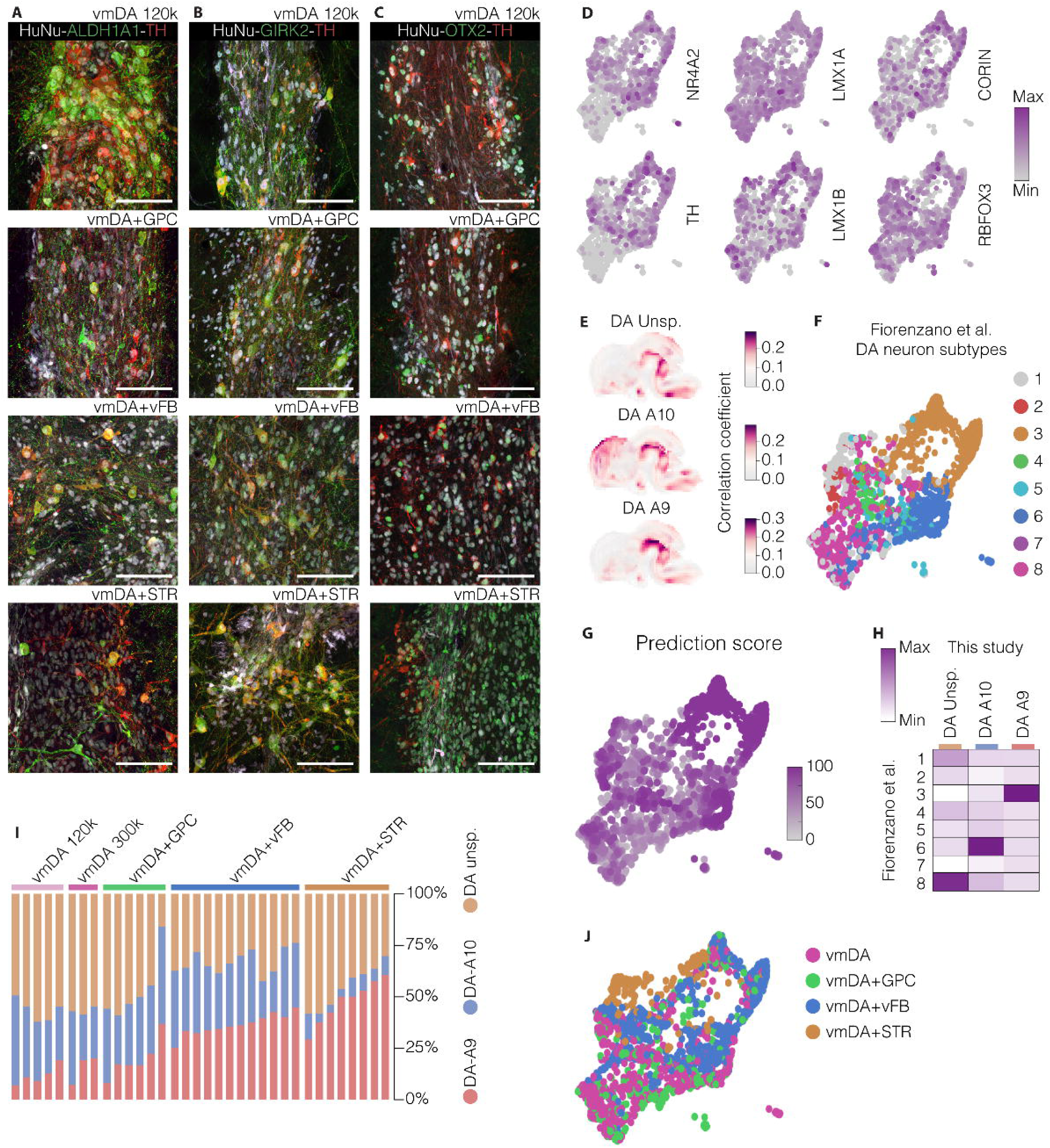
Characterization of vmDA-derived DA neurons in co-grafts. **A-C)** Immunohistochemistry for **A)** ALDH1A1, **B)** GIRK2 and **C)** OTX2 in combination with HuNu and TH in vmDA-only grafts and co-grafts. Scale bars = 100 μm. **D)** Feature plots showing the expression of selected midbrain and DA neuron markers in vmDA-derived DA neurons 28 weeks after transplantation. **E)** Correlation between the DA neuron transcriptome and spatial transcriptomics data from the postnatal day 4 (P4) mouse brain, generated using the VoxHunt package. **F-H)** Label transfer of DA neuron subtypes defined in Fiorenzano et al. 2024^19^. **F)**, including the relative prediction scores **G)** and a heatmap highlighting the correlation between DA subtypes 3, 6 and 8 and the A9-like, A10-like and unspecified clusters identified in this study **H)**. **I,J)** Barchart **I)** and SPRING plot **J)** showing the subtype composition of DA neurons across experimental groups and samples.

**Supplementary Figure 5.**
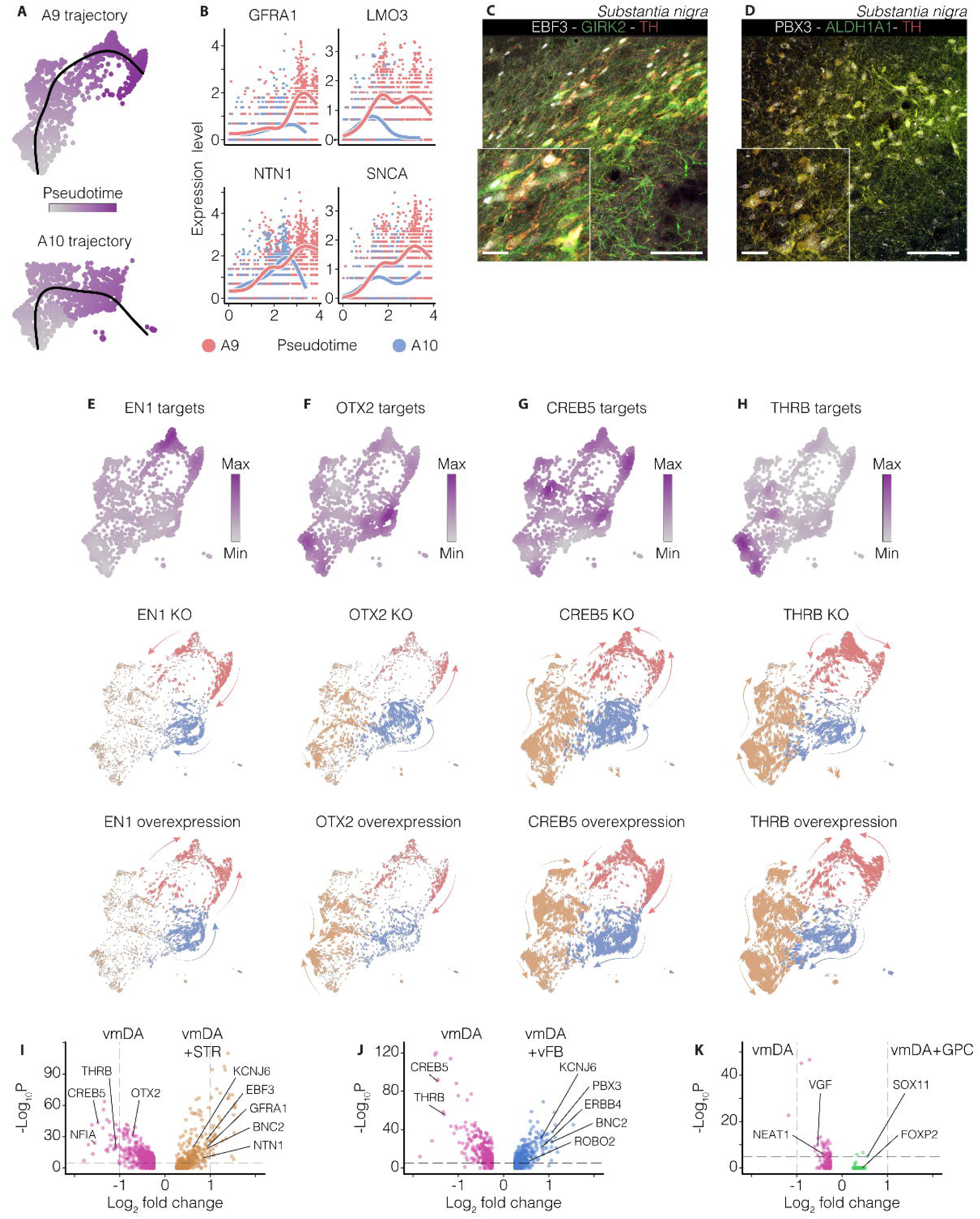
Trajectory analysis and transcriptional regulation of A9- and A10-like DA neurons. **A)** A9- and A10-like DA neuron trajectories inferred by slingshot on SPRING embeddings, with cells colour-graded based on pseudotime. **B)** Expression dynamics of selected markers along the pseudo-temporal axis, highlighting divergent transcriptional programs along the identified A9/A10 trajectories. **C,D)** Immunofluorescence staining for **C)** EBF3/GIRK2/TH and **D)** HuNu/PBX3/ALDH1A1 in the rat substantia nigra. Scale bars = 100 μm; inset 25 μm. **E-H)** Composite expression patterns of known target genes (top), and predicted effects of *in silico* transcription factor KO (middle) or overexpression (bottom) for EN1 **E)**, OTX2 **F)**, CREB5 **G)**, and THRB **H)**. **I-K)** Volcano plots showing differentially expressed genes between DA neurons derived from vmDA-only grafts and vmDA+STR co-grafts **I)**, or vmDA+vFB co-grafts **J)**, or vmDA+GPC co-grafts **K)**.

**Supplementary Figure 6.**
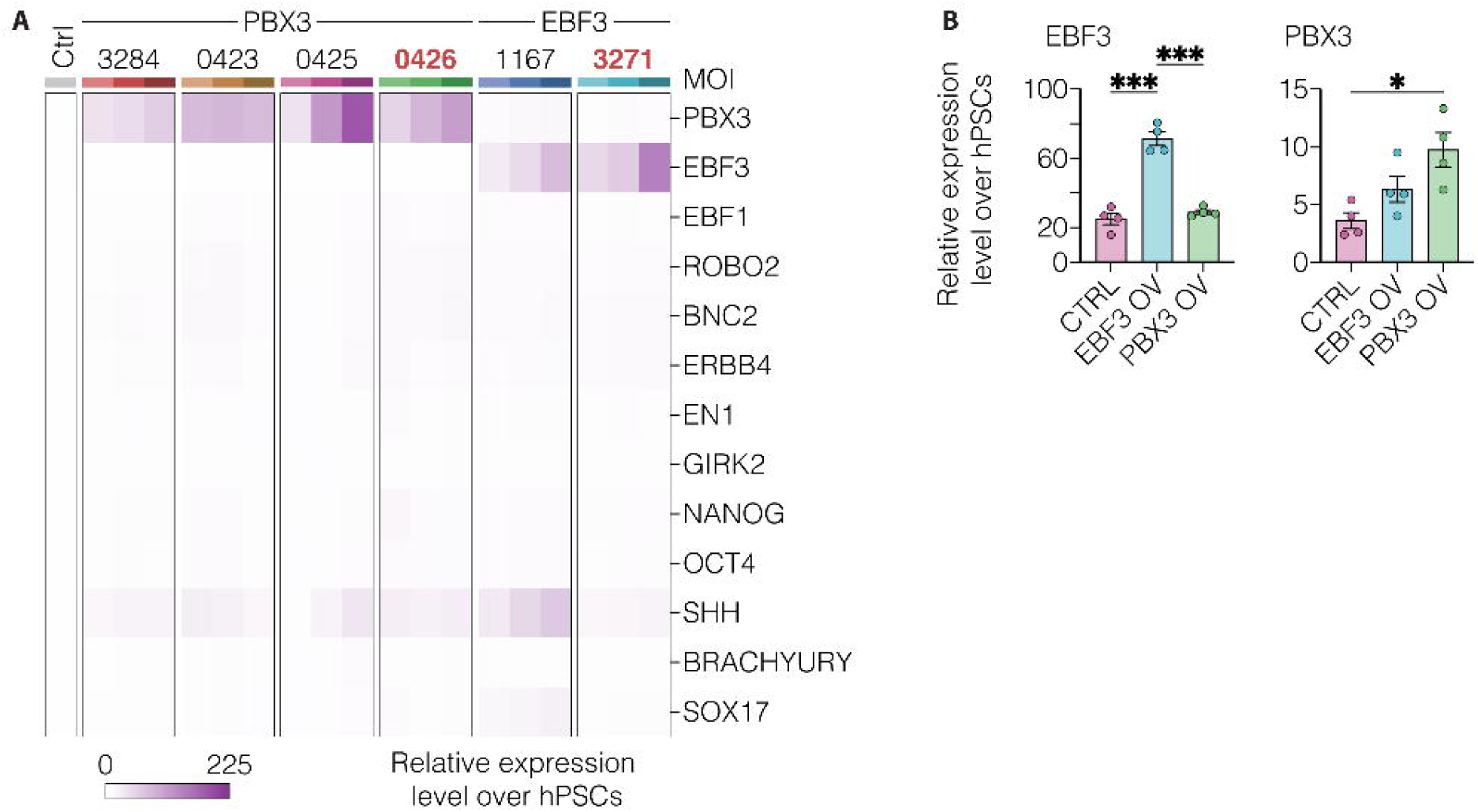
Selection and validation of ORFs for targeted *EBF3* and *PBX3* overexpression. **A)** Heatmap showing the average expression of *EBF3*, *PBX3*, selected downstream target genes, and lineage-specific markers in hPSCs overexpressing different vector sets^62^. For each ORF, MOIs of 1, 2, and 3 are shown from left to right. Vectors 0426 (*PBX3*) and 3271 (*EBF3*), selected for VM organoid experiments, were chosen based on robust induction of the intended target gene without detectable off-target effects on the other genes examined. **B)** Relative expression levels of *EBF3* and *PBX3* in VM organoids transduced with the two overexpression vectors at day 90.

